# The Rab GTPase Ypt1p governs the activation of Unfolded Protein Response (UPR) in *Saccharomyces cerevisiae* by promoting the preferential nuclear degradation of pre-*HAC1* mRNA

**DOI:** 10.1101/2022.08.18.504421

**Authors:** Sunirmal Paira, Anish Chakraborty, Biswadip Das

**Affiliations:** Department of Life Science and Biotechnology, Jadavpur University, Kolkata, India.

**Keywords:** Nuclear Exosome, CTEXT, UPR, Ypt1p, *RRP6*, *CBC1*, *HAC1*.

## Abstract

Induction of unfolded protein response (UPR) involves activation of transcription factor Hac1p that facilitates the transactivation of genes encoding ER-chaperones. Hac1p is encoded by *HAC1* pre-mRNA harboring an intron and a bipartite element (BE) at its 3′-UTR. This precursor RNA undergoes a reversible and differential intra-nuclear mRNA decay by the nuclear exosome/CTEXT at various phases of UPR. In this investigation, using a combination of genetic, and biochemical approach, the Rab-GTPase Ypt1p is demonstrated to control UPR signaling dynamics. Regulation of UPR by Ypt1p relies on its characteristic nuclear localization in absence of ER-stress resulting in its strong association with pre-*HAC1* mRNA at its 3′-UTR that promotes sequential recruitments of Nrd1-Nab3p-Sen1p (NNS) complex → CTEXT → the nuclear exosome onto the pre-*HAC1* mRNA that is accompanied by its rapid and selective nuclear decay. This accelerated 3′→5′ mRNA decay produces a pre-*HAC1* mRNA pool lacking the functional BE thus causing its inefficient targeting to Ire1p foci leading to their diminished splicing and translation. ER stress triggers a rapid relocalization of Ypt1p to the cytoplasm with its consequent dissociation from pre-*HAC1* mRNA thereby causing a decreased recruitment of NNS/exosome/CTEXT to precursor *HAC1* RNA leading to its diminished 3′→5′ degradation by the exosome. This diminished decay produces an increased abundance of pre-*HAC1* mRNA population with intact functional BE leading to its enhanced recruitment to Ire1p foci that is followed by its increased splicing and translation. This enhanced translation produces a huge burst of Hac1p that rapidly transactivates the genes encoding ER-chaperones.

## INTRODUCTION

In the nucleus of *Saccharomyces cerevisiae*, the nuclear exosome and its two co-factors, TRAMP and CTEXT selectively degrade a diverse spectrum of aberrant messages and thereby safeguards the cells from their detrimental effects (1–5). The nuclear RNA exosome is a large multi-protein complex of eleven subunits (1, 2, 6–11) and a few associated nuclear and cytoplasmic co-factors. Nine of these eleven subunits are arranged in a barrel (known as EXO9) within a core structure. The core structure consists of two stacked ring structures: a “trimeric” cap structure (composed of Rrp4p Rrp40p and Csl4p) is placed on the top of a “hexameric” ring (composed of Rrp41p, Rrp42p, Rrp43, Rrp45p, Rrp46p, and Mtr3p) with a central channel. This EXO9 structure contacts two catalytically active tenth and eleventh subunits, Dis3p/Rrp44p (possess both endo- and 3′→5′ exoribonuclease activity) and Rrp6p (3′→5′ exoribonuclease), respectively from opposite sides to form EXO11^Dis3p+Rrp6p^. Notably, a set of ancillary co-factors aids the EXO11^Dis3p+Rrp6p^ to selectively target and degrade distinct classes of target RNAs (12–14). TRAMP (**<u>TR</u>**f4/5p-**<u>A</u>**ir1/2p-**<u>M</u>**tr4p **<u>P</u>**olyadenylation) complex in *S. cerevisiae*, provides the best-characterized example of such exosome targeting complex, which consists of a non-canonical poly(A) polymerase, Trf4p/Trf5p, a DExH box RNA helicase, Mtr4p, and the Zn-knuckle RNA binding proteins, Air1p/2p (6, 15, 16). CTEXT (**<u>C</u>**bc1p-**<u>T</u>**if4631p-dependent **<u>EX</u>**osomal **<u>T</u>**argeting) complex (previously dubbed DRN for **<u>D</u>**ecay of **<u>R</u>**NA in the **<u>N</u>**ucleus) defines a second co-factor that consists of the nuclear mRNA-cap binding protein, Cbc1p/2p (17, 18), cytoplasmic non-sense mediated decay factor Upf3p (18, 19), translation elongation factor gamma, Tif4631p (18, 19) and a DEAD-box RNA helicase, Dbp2p. While TRAMP helps the nuclear exosome to target the aberrant messages generated in the early phase of mRNP biogenesis events, the CTEXT assists the exosome to target export-defective messages typically produced at the late phase of mRNA processing events and a small group of normal messages (19) called special mRNAs (20). A third protein complex, NNS (Nrd1p-Nab3p-Sen1p), previously known to participate in the transcription termination of sn- and snoRNA species (21–25) also plays a crucial role in the nuclear mRNA surveillance by acting as ‘exosome-specificity-factor’ (ESF) (26). NNS complex undergoes selective recruitment onto the aberrant/special messages co-transcriptionally by RNA polymerase II thereby ‘marking’ them as the target for the nuclear exosome (26).

A scrutiny of the stability of the global messages in WT, *rrp6-*Δ, and *cbc1-*Δ yeast strains by microarray analysis unveiled an enhancement of stability and steady-state levels of nearly two hundred normal messages including *HAC1* mRNA in both nuclear exosome-deficient *rrp6*-Δ and CTEXT-deficient *cbc1*-Δ strains (20). Notably, Hac1p is a b*ZIP* class of transcription factor which plays a pivotal role in the activation of Unfolded Protein Response (UPR) in *Saccharomyces cerevisiae* by trans activating the genes encoding ER chaperones (27–30). Remarkably, the precursor *HAC1* mRNA harboring an intron undergoes a reversible non-canonical splicing and subsequent translation to encode Hac1p. In the absence of stress, the precursor *HAC1* mRNA remains transnationally inactive via the formation of a secondary structural bridge between the intron and 5′-UTR (27). ER stress removes the intron via the non-spliceosomal splicing that is facilitated by the ER-resident kinase-endoribonuclease Ire1p and tRNA ligase Rlg1p thereby reinstating the translation of mature *HAC1* mRNA to produce an enormous amount of Hac1p (27–30). A crucial *cis*-acting bipartite element (BE) in the 3′-UTR of *HAC1* pre-mRNA/mRNA plays an instrumental role in recruiting/targeting this RNA to the specific intracellular foci consisting of active Ire1p, which rapidly cluster in response to stress (31).

Apart from the non-spliceosomal splicing, the *HAC1* pre-mRNA also undergoes another layer of regulation that involves a dynamic and reversible mRNA decay by the nuclear exosome/CTEXT in absence of ER-stress that resulted in a pool of precursor messages with a heterogeneous 3′-termini, most of which lack the functional BE. This pre-*HAC1* transcript pool fails to be targeted to the Ire1p foci in the absence of ER stress (32). ER stress triggers a diminution of the 3′→5′ exosomal decay of the pre-*HAC1* transcript body thereby producing an optimum repertoire of precursor *HAC1* messages, the majority of which harbor a functional and intact BE. Subsequently, this population of pre-*HAC1* mRNA with an intact/functional BE undergoes a more efficient targeting and recruitment to the Ire1p clusters (32). The nuclear exosome/CTEXT, thus, exert a fine-tuning in the regulation of UPR signaling dynamics by facilitating the production of the functional pre-*HAC1* mRNA population with an intact BE during the absence and presence of ER-stress (32). However, how the precursor *HAC1* mRNA is specifically recognized and targeted for exosomal decay currently remains obscure.

A previous investigation identified an small Rab-family GTPase Ypt1p as a top interactor of *HAC1* pre-mRNA (33). Ypt1p is an essential modulator of ER-to-Golgi transport in the secretory pathway (34). Using an *in vitro* proteomic approach, these researchers demonstrated that Ypt1p binds to *HAC1* pre-mRNA in a UPR-dependent manner, which facilitates its decay in the unstressed cells, and knocking down Ypt1p led to the elevated levels of *HAC1* pre-mRNA due to diminished decay (33). However, this study did not address the mechanism of Ypt1p-dependent decay of pre-*HAC1* mRNA. Since this essential Rab-GTPase interacts very strongly with the pre-*HAC1* mRNA in absence of ER stress, we addressed if Ypt1p promotes the accelerated decay of the pre-*HAC1* message. Using a combination of genetic, biochemical, and cell-biological approaches we demonstrate that Ypt1p indeed coordinates the sequential recruitment of the major ESF NNS, CTEXT, and finally, the nuclear exosome to promote the preferential degradation of the precursor *HAC1* mRNA and thereby limits its transport and recruitment to Ire1p. ER stress promotes nucleus-to-cytoplasmic relocalization of Ypt1p resulting in its reduced association with pre-*HAC1* mRNA leading to diminished exosomal decay, increased targeting and delivery of this precursor RNA to Ire1p clusters followed by enhanced splicing and translation. This augmented translation finally results in the huge burst of Hac1p abundance, which rapidly trans-activates genes encoding the ER chaperone.

## RESULTS

### Ypt1p tunes the unfolded protein response (UPR) signaling dynamics in *Saccharomyces cerevisiae*

Although the previous investigation displayed an association of Ypt1p with unspliced *HAC1* mRNA in absence of ER-stress and thereby regulates its steady-state levels and stability (33), the ability of the *ypt1* mutant cells to grow and survive in the absence and presence of ER stress was not studied. We initiated this investigation by verifying if the inactivation of Ypt1p influences the ability of the yeast cells to grow and survive in presence of ER stress. To this end, we examined the growth profiles of WT and *ypt1-3* mutant yeast strains in the absence and presence of either 1μg/ml Tunicamycin (Tm) (experiments conducted in both solid and liquid medium) or 5 mM DTT (experiments carried out in liquid medium) (Fig 1A-C). As shown in these figures, in all cases mutant *ypt1-3* strain displayed a better and more efficient growth in presence of both Tunicamycin and DTT (Fig 1A-C). These findings are reminiscent of our previous observation that inactivation of either the nuclear exosome component Rrp6p or CTEXT component Cbc1p led to the increased sustenance of *cbc1-*Δ and *rrp6-*Δ mutant yeast strains (32). To explore further if the inactivation of *YPT1* affects the cellular levels of the precursor/mature *HAC1* mRNA, we determined their steady-state levels (Fig 1D-E) and the stability (Fig. 1F). The results of these experiments revealed that indeed the abundance of both the mature and pre-*HAC1* mRNA populations increased by about 1.5 and 2 folds respectively in absence of ER-stress in the *ypt1-3* mutant strain. Importantly, the level of the *HAC1* pre-mRNA displayed a significant (two folds) enhancement in a selective fashion in the *ypt1-3* mutant strain in absence of tunicamycin (Fig. 1E, lower histogram). Exposure of these strains to ER stress led to an even further increment in the level of the pre-*HAC1* mRNA (≈ 3-3.5 folds) in *ypt1-3* strain relative to WT in unstressed condition (Fig. 1E, lower histogram). Consistent with this finding, a comparison of the decay rate and half-life of pre-*HAC1* mRNA between WT and *ypt1-3* strains displayed that the decay is diminished in the *ypt1-3* strain with a concomitant enhancement of its stability by about 2.5 folds (WT HL=10 minutes, *ypt1-3* HL=27 minutes) (Fig. 1E). This data thus strongly indicates that the difference in the steady-state levels of pre-*HAC1* mRNA as observed in Fig. 1D-E is correlative to its diminished decay rate in *ypt1-3* yeast strain. Interestingly, no significant alterations in either the steady-state levels or decay rates of three non-ER-specific mRNAs, *CYC1*, *LYS2,* and *NCW2* were noted in the *ypt1-3* strain relative to WT (Supplementary Fig S1A-B). These findings collectively suggest that Ypt1p indeed plays a vital role in modulating the UPR signalling dynamics by facilitating the decay of the pre-*HAC1* mRNA, which is specific and physiologically relevant thereby validating the findings from the previous report (33).

**Figure 1.**
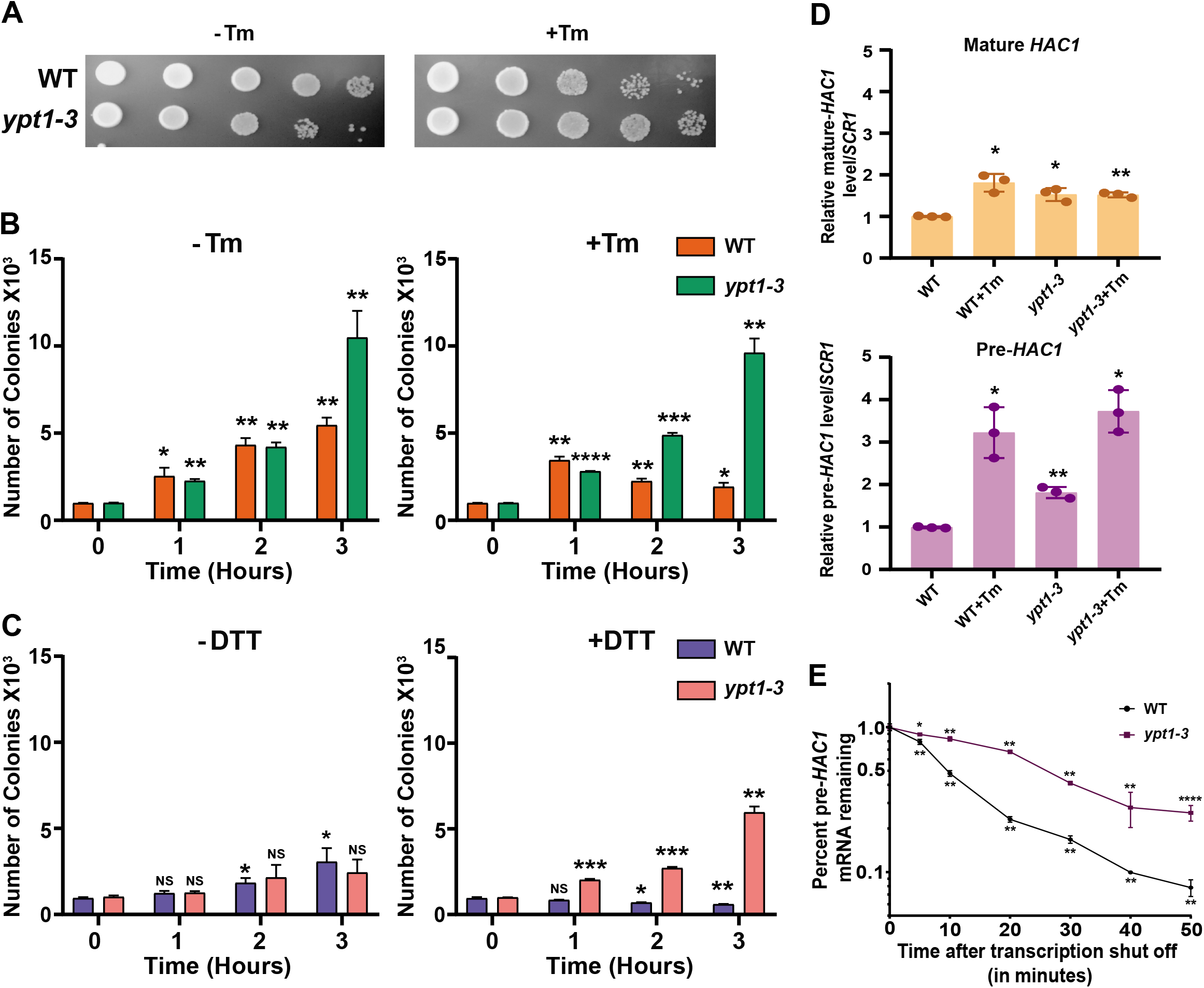
Ypt1p plays a vital role in the activation of UPR in baker’s yeast: **A.** Relative growth of WT (yBD-433), and *ypt1-3* (yBD-434) strains in YPD solid growth media in absence (-Tm) and in presence of 1 µg/ml tunicamycin (+Tm). Equal number of cells of WT and *ypt1-3* yeast strains grown in YPD liquid medium in absence of tunicamycin were spotted with ten-fold dilutions on solid YPD medium without or with 1 µg/ml tunicamycin and grown at 30°C for 72 hours before photographed. **B.** Relative growth of WT (yBD-433) (orange Bar) and *ypt1-3* (yBD-434) (green Bar) strains in YPD liquid growth media either in absence (-Tm) or in presence of 1 µg/ml tunicamycin (+Tm). Each of these strains was grown in liquid YPD medium in absence of tunicamycin to an optical density of 0.6 and was divided into two parts. In one-half of the culture, tunicamycin was added to a final concentration to 1 µg/ml (+Tm) whereas equal volume of DMSO was added to other half of the culture (-Tm). Both cultures were then continued to grow for 3 hours and 100 µl of aliquots of each culture was withdrawn at indicated times after tunicamycin (or DMSO) addition, washed once to remove the drug/DMSO and spreaded on the solid YPD plate. Plates were incubated for 72 hours at 30°C and the colony numbers were scored and plotted as histograms which represents the mean of three independent experiments (n=3) and the standard error of means (error bars). **C**. Relative growth of WT (yBD-433) (blue Bar) and *ypt1-3* (yBD-434) (pink Bar) strains in YPD liquid growth media in absence (-DTT) and in presence of 5 mM Dithiothreitol (+DTT). WT and *ypt1-3* strains were grown in liquid YPD medium in absence of DTT to an optical density of 0.6 and were divided into two parts. In one-half of the culture, DTT was added to a final concentration to 5 mM (+DTT) whereas equal volume of DMSO was added to other half of the culture (-DTT). Cultures were continued to grow for an additional 3 hours and 100 µl of aliquots were withdrawn at indicated times after DTT addition and were spreaded onto solid YPD plate. Plates were incubated for 72 hours at 30°C and the colony numbers were scored and plotted as histograms which represents the mean of three independent experiments (n=3) and the standard error of means (error bars). **D.** Histogram depicting the steady state levels of only mature (upper panel) and precursor *HAC1* (lower panel) mRNA levels in WT (yBD-433) and *ypt1-3* (yBD-434) strains in the absence and presence of tunicamycin as determined by qPCR analysis. Transcript copy numbers/2ng cDNA of each strain was normalized to *SCR1* RNA levels in respective strains and are presented as means ± SE (n=3 for each strain). Normalized value of individual mRNA from WT strain was set to one. **F.** Decay rates of pre-*HAC1* mRNA in WT (black line) and *ypt1-3* (violet line) as determined by qRT-PCR. Decay rates were determined from three independent experiments (biological replicates) by qRT-PCR analysis (using primer sets of *HAC1* intronic sequences) and the intronic signals were normalized to *SCR1* RNA signals obtained for each time point and the normalized signals (mean values ± SD) were presented as the fraction of remaining pre-*HAC1* RNA (with respect to normalized signals at 0 min) as a function of time after shutting-off the transcription of RNA polymerase II by the addition of 1, 10-phenanthroline. For panels D and E, the statistical significance of difference as reflected in the ranges of p-values estimated from Student’s two-tailed t-tests for a given pair of test strains for each message are presented with following symbols, ∗<0.05, ∗∗<0.005 and ∗∗∗<0.001, NS, not significant.

### Ypt1p promotes the nuclear decay of pre-*HAC1* mRNA by facilitating the recruitment/ targeting of the Nuclear Exosome and the CTEXT Complex to the premature *HAC1* mRNA

Despite the recognized role of Ypt1p in binding the unspliced *HAC1* mRNA and regulating the UPR signalling dynamics (33), its underlying mechanism is currently unknown. Notably, unspliced pre-*HAC1* mRNA undergoes a dynamic and reversible degradation in the nucleus by the nuclear exosome and its co-factor CTEXT (Sarkar et al., 2018). These findings let us postulate that Ypt1p facilitates the recruitment of the nuclear exosome and CTEXT to the pre-*HAC1* mRNA during the unstressed condition to promote its accelerated decay. To this end, we first query if Ypt1p interacts with the nuclear exosome/CTEXT by carrying out an immuno-precipitation experiment by anti-TAP antibody followed by the detection of exosome/CTEXT components using their specific antibodies using the cell extracts prepared from WT strain expressing Ypt1p-TAP grown either in the absence or presence of tunicamycin. As shown in Fig. 2A, indeed Ypt1p displays strong interactions with the nuclear exosome component, Rrp6p/Rrp4p and CTEXT components, Cbc1p/Tif4631p (Fig. 2A, Lane 3, supplementary figure Fig. S1C) in absence of ER-stress. Interestingly, interactions of Ypt1p with all of the Rrp6p, Rrp4p, Cbc1p, and Tif4631p reduced dramatically when ER stress was imposed (Fig. 2A, Lane 2). Importantly, mutant Ypt1-3p did not exhibit a similar interaction with any of these exosome/CTEXT components either in the absence or presence of ER stress in a similar immuno-precipitation experiment that was carried out with the extract prepared from the strain expressing the TAP-tagged version of the mutant form of Ypt1p, Ypt1-3p-TAP (Fig. 2B, lanes 2 and 3). Findings from these experiments suggest that in absence of ER stress Ypt1p indeed interacts strongly with the exosome and CTEXT components, which was reversed when ER stress was employed.

**Figure 2:**
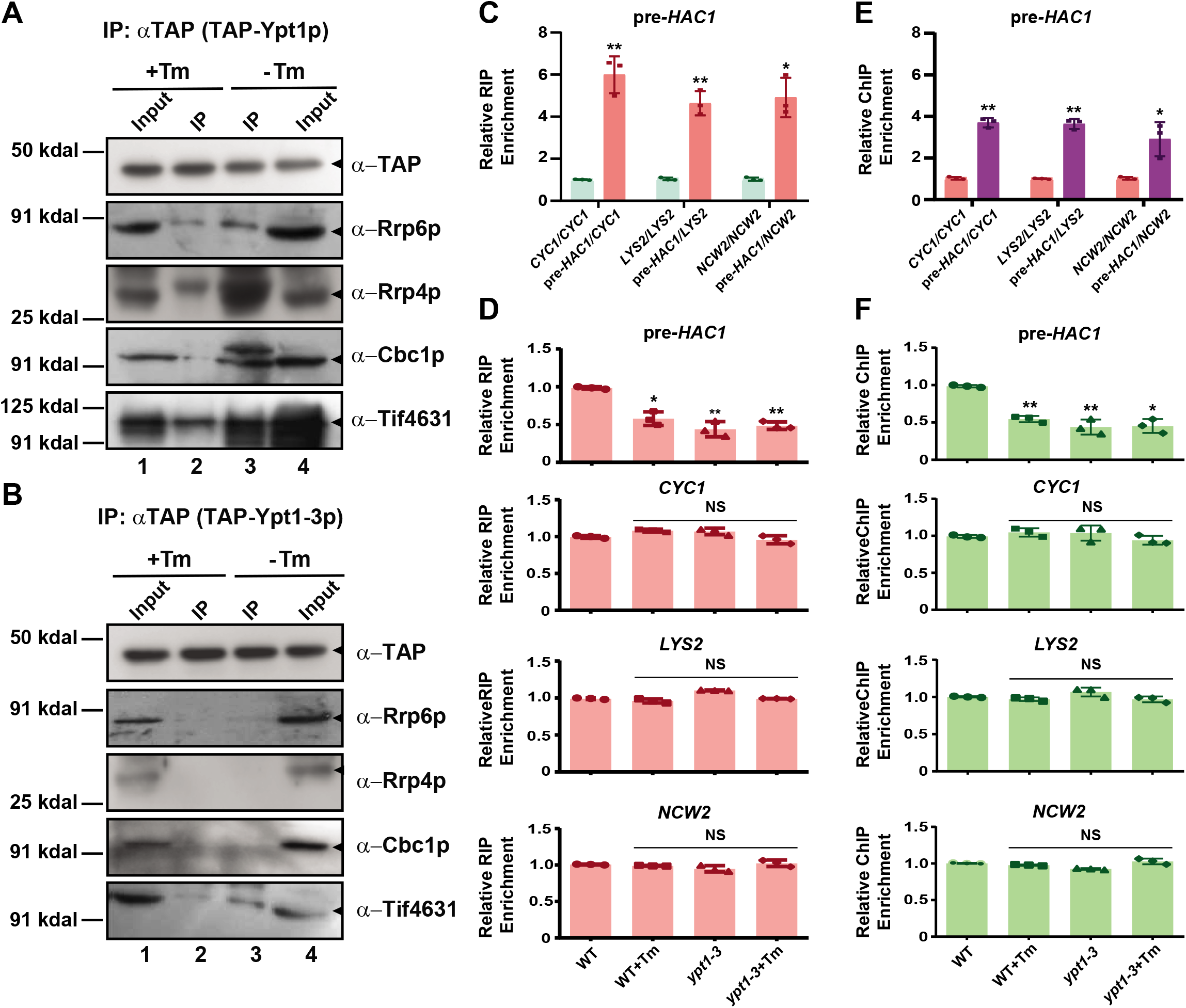
Ypt1p promotes the decay of precursor *HAC1* mRNA by promoting the co-transcriptional recruitment of the nuclear exosome and CTEXT onto it. **A-B** Immunoblots depicting the reversible physical association of Ypt1p with the nuclear exosome components, Rrp6p and Rrp4p and CTEXT components, Cbc1p and Tif4631p. Yeast cells expressing either WT Ypt1p-TAP **(A)** or mutant Ypt1-3p-TAP **(B)** were grown in the absence and in the presence of tunicamycin (1 μg/ml). Cells were subsequently harvested and Ypt1p was precipitated by standard immunoprecipitation as described in **Materials and Methods** followed by the detection of Rrp6p, Rrp4p, Cbc1p, and Tif4631p using their respective antibodies. The position of the nearest and relevant MW marker (in kDal) in each western blot panel is indicated at the left. **C.** Relative efficiency of binding of Ypt1p-TAP to the pre-*HAC1* mRNA relative to its binding to different non UPR-specific mRNAs in unstressed condition as determined by quantification of bound pre-*HAC1* and other non UPR-specific RNAs recovered from IP following standard immunoprecipitation with anti-TAP antibody using the extract from yeast strain expressing Ypt1p-TAP. The ratio (mean±SE) of abundance of pre-*HAC1* recovered from the IP relative to that of a non ER-specific RNA is presented as Relative enrichment. In each case, the abundance of a non-UPR specific RNA was set to one. **D.** Ypt1p strongly and selectively binds to the *HAC1* pre-mRNA in the absence of ER stress and ER-stress weakens Ypt1p-pre-*HAC1* interaction. Cells expressing either WT Ypt1p-TAP (yBD-457) or mutant Ypt1-3-TAP (yBD-565) were grown in the absence or presence of tunicamycin (1µg/ml) followed by the exposure of the culture to UV in order to induce RNA-protein cross-linking as described in **Materials and Methods**. After harvesting, the Ypt1p and Ypt1-3p was precipitated by anti-TAP antibody followed by the extraction of RNA from the IP and detection of either pre-*HAC1* mRNA or a series of non UPR-specific *CYC1*, *LYS2* and *NCW2* mRNAs by qRT-PCR. **E.** Relative co-transcriptional recruitment of Ypt1p onto pre-*HAC1* mRNA relative to its recruitment onto other non UPR-specific mRNAs as determined by RNA Chromatin Immunoprcipitation (RNA-ChIP) using TAP antibody as described in **Materials and Methods**. In each case, the co-transcriptional recruitment of a non-UPR specific RNA was set to one and the relative recruitment of pre-*HAC1* mRNA is expressed as retive enrichment relative to the non-specific *CYC1*, *LYS2* and *NCW2* mRNAs. **F**. Ypt1p selectively recruited co-transcriptionally onto pre-*HAC1* mRNA in the absence of ER stress and ER stress reverses the recruitment. Yeast cells expressing WT Ypt1p-TAP (yBD-457) and mutant Ypt1-3-TAP (yBD-565) were grown in the absence or presence of tunicamycin (1µg/ml final concentration) followed by the addition of 5% formaldehyde to the culture to induce chromatin-RNA-protein cross-linking. After harvesting, the Ypt1p and Ypt1-3p was precipitated by anti-TAP antibody followed by the de-crosslinking of DNA and detection of either pre-*HAC1* mRNA or non-UPR specific *CYC1*, *LYS2* and *NCW2* mRNAs by qRT-PCR. Their abundance was presented as histogram/scattered plot (mean±SE) that expresses the relative Ypt1p ChIP enrichment. For both the RIP and RNA-ChIP experiments, the average of three independent experiments is shown where immunoprecipitated (output) samples were normalized to input following quantification by quantitative real-time PCR. The error bars in the graph represent standard deviations. The statistical significance of difference as reflected in the ranges of p-values were estimated from Student’s two-tailed t-tests for a given pair of test strains for each message are presented with following symbols, * <0.05,**<0.005 and ***<0.001, NS, not significant.

Next, we assessed if this physical association of Ypt1p with the nuclear exosome/CTEXT is physiologically meaningful and to this end, we checked if Ypt1p (i) interacts with pre-*HAC1* mRNA *in vivo* in the presence and absence of ER stress and (ii) is recruited specifically onto the pre-*HAC1* mRNA in a co-transcriptional manner. To address the first query, we carried out an RNA-immuno-precipitation (RIP) experiment from the UV-cross-linked extracts prepared from yeast strains expressing Ypt1p-TAP with an anti-TAP antibody followed by the estimation of the recovered pre-*HAC1* mRNA in the immunoprecipitate (IP) using qRT-PCR (Fig. 2C). As shown in this figure, the abundance of the pre-*HAC1* mRNA recovered from the Ypt1p-TAP IP displayed a ≈ 5-6 folds enrichment relative to three non-ER-specific mRNAs (not associated with UPR signalling), *CYC1*, *LYS2* and *NCW2* that could be retrieved from the same IP sample (Fig. 2C) thereby demonstrating the specificity of the Ypt1p-*HAC1* pre-mRNA interaction. Furthermore, a comparison of pre-*HAC1* mRNA bound to Ypt1p in the absence and presence of ER-stress showed that the abundance of the recovered pre-*HAC1* from Ypt1p-TAP IP significantly diminished in presence of ER-stress relative to unstressed WT cells (Fig. 2D, top histogram). As predicted, the levels of bound pre-*HAC1* mRNA that were recovered from Ypt1-3-TAP IP were significantly lower than that recovered from the functional Ypt1p (Fig. 2D, top histogram) suggesting that the Ypt1p-pre-*HAC1* mRNA interaction diminished dramatically in a yeast strain carrying the *ypt1-3* allele. No significant alteration of the recovered *HAC1* pre-mRNA was noted in *ypt1-3* strains when ER stress was imposed (Fig. 2D, top histogram). In addition, no significant strain-specific (WT vs. *ypt1-3* mutant) or ER-stress-specific (without vs. with ER-stress) differences were noted in the abundance of the recovered non-ER-specific *CYC1*, *LYS2,* and *NCW2* messages (Fig. 2D, three bottom histograms). This finding indicates a strong and specific interaction between Ypt1p to pre-*HAC1* mRNA in absence of ER stress (Fig. 2C and D), which was significantly reversed when ER stress was applied.

To further query if this binding/interaction between Ypt1p and pre-*HAC1* mRNA is forming during its nuclear transcription, we employed the chromatin-immuno-precipitation (ChIP) assay. It should be noted here that, although the ChIP assay was believed to crosslink chromatin/DNA with the DNA-binding proteins, a vast majority of RNA binding proteins also remain associated with the transcribing messenger RNAs, which remain physically associated with the chromatin. RNA-ChIP, adapted from the classical DNA-ChIP, is now widely used to study and analyze the binding profiles of transcription, splicing, export, and mRNA decay factors, which are recruited co-transcriptionally and remained strongly associated with the transcribing/maturing nascent mRNAs (26, 32, 35, 36). Before carrying out the actual comparison of Ypt1p occupancy profiles onto pre-*HAC1* mRNA in various yeast strains during the ER-stress conditions by RNA-ChIP, we first determined if the ChIP signal generated from our assay procedure is highly specific. In this experiment, we used the fragmented chromatin prepared from yeast cells expressing Ypt1p-TAP that is precipitated with the anti-TAP antibody followed by quantification of the abundance of chromatin-bound pre-*HAC1* mRNA by qRT-PCR (Fig 2E). As shown in Fig. 2E, the abundance of the pre-*HAC1* mRNA recovered from the Ypt1p-TAP chromatin-IP samples exhibited a ≈ 3-4 folds enrichment relative to the abundance of the three other non-ER-specific mRNAs, *CYC1*, *LYS2* and *NCW2* that could be recovered from the same chromatin-IP samples (Fig. 2E). Subsequently, the abundance of pre-*HAC1* mRNA recovered from anti-TAP chromatin-IP from the WT yeast cells in absence of ER stress was estimated to be two-fold higher relative its abundance in presence of ER stress (Fig. 2F, top histogram). Moreover, the levels of *HAC1* mRNA bound to chromatin-IP done from the yeast cell expressing Ypt1-3p mutant protein with the anti-TAP antibody are significantly lower in the *ypt1-3* yeast strain relative to WT in absence of ER-stress and this level did not alter any further when this strain was challenged with ER-stress. As noted before, no significant strain-specific and ER-stress-specific alterations were observed in the binding profiles of the three non-ER-specific, *CYC1*, *LYS2,* and *NCW2* mRNAs thereby suggesting the specificity of Ypt1p-pre-*HAC1* mRNA interaction during its Ypt1p-dependent co-transcriptional recruitment. These findings, collectively suggest that a large fraction of Ypt1p is recruited onto pre-*HAC1* mRNA in absence of ER stress in a transcription-dependent manner, which is significantly reduced when WT yeast cells encounter ER stress.

### Ypt1p selectively recruits Nrd1p onto the pre-*HAC1* mRNA leading to the further recruitment of the nuclear exosome and the CTEXT

Recent observation from our lab suggests that during the nuclear mRNA surveillance, the trimeric Nrd1p-Nab3p-Sen1p (NNS) complex is preferentially loaded onto diverse sets of aberrant and special messages thereby ‘marking’ them as the exosomal targets (Singh et al., 2021). This observation prompted us to address whether Ypt1p recruits the exosome/CTEXT directly or via the intermediate loading of the NNS complex. Alternatively, the NNS complex may also be recruited onto the pre-*HAC1* mRNA before of the Ypt1p recruitment. To delineate this issue, we first examined if Ypt1p displays any physical association with the NNS component, Nrd1p, and Nab3p by co-immunoprecipitation with the anti-TAP antibody using the extract prepared from WT yeast strain expressing Ypt1p-TAP followed by the detection of Nrd1p and Nab3p using either anti-Nrd1p or anti-Nab3p antibodies. As shown in Fig 3A, Ypt1p displays a strong association with both NNS components, Nrd1p and Nab3p in absence of ER-stress (Fig. 3A, lane 2), which either became significantly decreased (Nrd1p) or completely abrogated (Nab3p) when ER-stress was employed (Fig. 3A, lane 2). Importantly, the mutant form of this protein Ypt1-3p did not exhibit any association with any of the NNS components, neither in the absence nor the presence of ER-stress (Fig. 3B). This data implies that possibly Ypt1p co-ordinates the recruitment of the nuclear exosome and CTEXT via the loading of the NNS complex in the intermediate stage. This hypothesis predicts that the NNS complex plays a vital role in the UPR signaling dynamics by recruiting the exosome/ CTEXT onto the pre-*HAC1* mRNA co-transcriptionally. Consequently, in an *nrd1-1* mutant yeast strain, the level of pre-*HAC1* mRNA would display a much higher level and a yeast strain carrying *nrd1-1* would exhibit better sustenance to ER stress. We verified the first prediction, by examining the steady-state levels of pre-*HAC1* mRNA in WT and *nrd1-1* strains by qRT-PCR assay, which demonstrated that indeed the pre-*HAC1* mRNA level was significantly increased in *nrd1-1* strain (Fig. 3B). To bolster this finding and validate the second speculation, we examined the growth profiles of WT and *nrd1-1* mutant yeast strains in the absence and presence of 1μg/ml Tunicamycin (supplementary Fig S2A). As shown in this figure, the mutant *nrd1-1* yeast strain displayed a better and more efficient growth in presence of ER stress (supplementary Fig S2A). These findings collectively affirm that the precursor *HAC1* RNA level was enhanced in the *nrd1-1* yeast strain due to its diminished decay leading to the increased sustenance of this mutant yeast strain to ER-stress, thereby supporting the idea that NNS complex plays a pivotal role in the UPR signalling dynamics in baker’s yeast.

**Figure 3:**
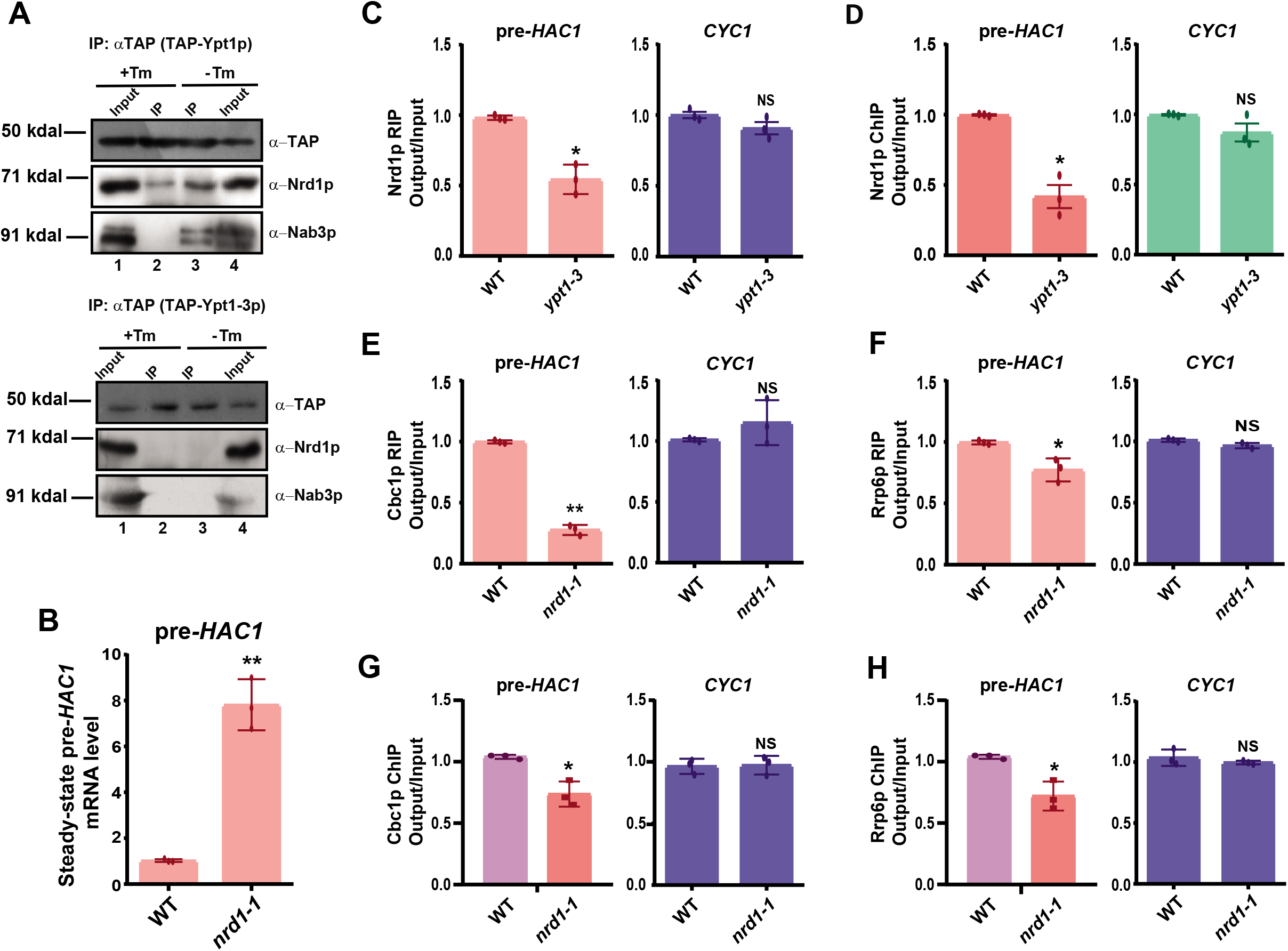
Ypt1p coordinates the co-transcriptional recruitment of NNS complex onto the precursor *HAC1* message in absence of ER-stress, which further stimulates the recruitment of the nuclear exosome and CTEXT on it. **A.** Immunoblots depicting the reversible physical association of Ypt1p with NNS components Nrd1p and Nab3p in the absence of ER stress. Yeast cells expressing either WT Ypt1p-TAP (yBD-457) or mutant Ypt1-3-TAP (yBD-565) were grown in the absence and presence of tunicamycin (1 µg/ml final concentration) followed by harvesting and preparation of cell extracts as described in **Materials and Methods**. Ypt1p was precipitated by standard immunoprecipitation followed by the detection of Nrd1p and Nab3p using their respective antibodies. The position of the nearest and relevant MW marker (in kDal) in each western blot panel is indicated at the left. **B.** Histogram depicting the steady state levels of pre-*HAC1* mRNA levels in WT (yBD-148) and *nrd1-1* (yBD-177) yeast strains using qPCR analysis using a primer pair specific to *HAC1* intron. Transcript copy numbers/2 ng cDNA of each strain was normalized to *SCR1* RNA levels in respective strains and are presented as means ± SE (n=3 for each strain). Normalized value of pre-*HAC1* mRNA from WT samples was set to one. **C-D.** Ypt1p dictates the specific binding (**C**) and co-transcriptional recruitment (**D**) of Nrd1p to pre-*HAC1* message in absence of ER-stress. Cells expressing either WT Ypt1p (yBD-433) or mutant Ypt1-3p (yBD-434) were grown in absence of ER-stress followed by either exposing the culture to UV to induce RNA-protein crosslinking (for RIP experiment presented in **C**) or treating the culture with 5% formaldehyde to induce chromatin-RNA-protein cross-linking (for RNA-ChIP experiment presented in **D**). After harvesting of the cells, Nrd1p was precipitated by anti-Nrd1 antibody that was followed either by the extraction of RNA and detection of pre-*HAC1* and *CYC1* mRNAs (**C**) or by the de-crosslinking of the chromatin-RNA and detection of pre-*HAC1* and *CYC1* mRNAs (**D**) by qRT-PCR. Their abundance were presented as scattered plot showing the mean and distribution of individual data points. **E-F**. Nrd1p dictates the specific binding of exosome and CTEXT components, Rrp6p and Cbc1p respectively to pre-*HAC1* message in absence of ER-stress. WT (yBD-157) and *nrd1-1* (yBD-158) yeast strains both expressing Rrp6p-TAP were grown in liquid YPD medium followed by UV-directed RNA-protein crosslinking before harvesting. Immunoprecipitation was subsequently performed using specific anti-Cbc1p **(E)** and anti-TAP antibody **(F)** followed by qRT-PCR analyses to quantify pre-*HAC1* and *CYC1* mRNAs. The average of the three independent experiments are shown where IP (output) samples were normalized to input signals following quantification by qRT-PCR. The error bars in the graph represent standard deviations. **G-H.** Nrd1p1p promotes the co-transcriptional recruitment of the CTEXT component Cbc1p (**G**) and exosome component Rrp6p (**H**) to pre-*HAC1* message in absence of ER-stress. WT (yBD-157) and *nrd1-1* (yBD-158) yeast strains both expressing Rrp6p-TAP were grown in liquid YPD medium followed by formaldehyde-directed chromatin-RNA-protein crosslinking before harvesting. Immunoprecipitation was subsequently performed using specific anti-Cbc1p and anti-TAP antibody followed by extraction of chromatin and quantification of pre-*HAC1* and *CYC1* mRNAs by qRT-PCR. The average of three independent experiments is shown where ChIP (output) samples were normalized to input following quantification by real-time PCR. In all cases, the normalized RIP or ChIP signals obtained in WT strain was set to one. The error bars in the graph represent standard deviations. The statistical significance of difference as reflected in the ranges of p-values estimated from Student’s two-tailed t-tests for a given pair of test strains for each message are presented with following symbols, * <0.05, **<0.005 and ***<0.001, NS, not significant.

In the next step, we verified that both the Nrd1p binding to pre-*HAC1* mRNA (determined by RIP) (Fig. 3C, left histogram) and its co-transcriptional recruitment (determined by ChIP) (Fig. 3D, left histogram) onto pre-*HAC1* mRNA became significantly diminished in the yeast strain carrying the *ypt1-3* allele. In contrast, the Ypt1p binding (determined from Ypt1p RIP) and recruitment (determined from Ypt1p ChIP) did not alter in the *nrd1-1* strain relative to the WT strain (supplementary Fig. S2D-E). This finding strongly argued that although Nrd1p recruitment onto pre-*HAC1* mRNA is Ypt1p-dependent, Ypt1p recruitment is independent of the presence of functional Nrd1p. Collective data thus supports the idea that Ypt1p is recruited onto pre-*HAC1* mRNA upstream of Nrd1p (NNS complex), which in turn further recruits NNS complex onto it. Further, we query if the binding (Fig. 3E-F) and the co-transcriptional recruitments (Fig. 3G-H) of the exosome component Rrp6p and CTEXT component Cbc1p to the pre-*HAC1* mRNA requires functional Nrd1p. The results of these experiments demonstrated that the binding (assessed by their RIP signals) and co-transcriptional recruitment (assessed by their ChIP signals) of Rrp6p and Cbc1p onto pre-*HAC1* mRNA are significantly impaired in yeast strain harboring *nrd1-1* allele relative to WT strain (Fig. 3E-H, left histogram in each case). The relative binding and recruitment of these decay factors on another non-ER specific CYC1 mRNA did not alter under any of these conditions (Fig. 3E-H, right histogram in each case). Collectively, these data are supportive of the view that the Rab-GTPase, Ypt1p after being recruited onto the *HAC1* pre-mRNA in a co-transcriptional manner co-ordinates further recruitment of the NNS complex. Recruitment of NNS complex is followed by further co-transcriptional recruitment of the nuclear exosome component Rrp6p and CTEXT component, Cbc1p on to the *HAC1* pre-mRNA thereby promoting its rapid nuclear decay in absence of ER-stress.

### Ypt1p promotes the targeting and recruitment of the pre-*HAC1* mRNA to the Ire1 foci

The extent of the nuclear exosome/CTEXT-dependent kinetic/reversible nuclear degradation of the precursor *HAC1* mRNA determines the fraction of the pre-*HAC1* message population harboring the bipartite-element (BE) at its 3′-UTR generated under various stress conditions (32). The experiments presented above revealed that the dynamic and reversible nuclear decay of *HAC1* pre-mRNA is primarily dictated by the reversible and regulated recruitment/loading of Rab-GTPase Ypt1p on the unspliced *HAC1* mRNA. Ypt1p recruitment further promotes its decay by sequential downstream recruitment of the NNS, the nuclear exosome, and CTEXT. These data, therefore, encouraged us to interrogate if Ypt1p governs the targeting of the pre-*HAC1* mRNA from its transcription site and influences its successful recruitment to the Ire1p foci. We validated this query by determining the intracellular distribution of *HAC1* mRNA in WT and *ypt1-3* yeast strains. Both of these strains co-express *HAC1^NRE^* mRNA (an engineered version of *HAC1* construct carrying an RNA aptamer with a 22 nt RNA module identical to nucleolin recognition element, NRE), a Nucleolin-GFP fusion protein, and Ire1p-RFP (Ire1p protein tagged with RFP) (37) using confocal imaging. The *in situ* distribution/localization of pre-*HAC1^NRE^* RNA bound Nucleolin-GFP (*HAC1^NRE^*-Nucleolin-GFP) fusion protein, and Ire1p-RFP (Fig. 4A, see merge and DIC/merge panel in the top row) clearly shows that in WT cells, where Ypt1p is functioning optimally, GFP signal representing the *HAC1* message appears to be excluded from the red spots representing localized Ire1p clusters. In the *ypt1-3* mutant strain, in contrast, a comparison of the distribution of pre-*HAC1^NRE^* bound Nucleolin-GFP and Ire1p-RFP revealed the presence of more abundant yellow spots that resulted from the overlapping GFP and RFP signals (Fig. 4A, see merge and DIC/merge panel in bottom row). This finding thus indicates that the pre-*HAC1* message is more efficiently targeted to the Ire1p foci in the *ypt1-3* mutant yeast strain.

**Figure 4:**
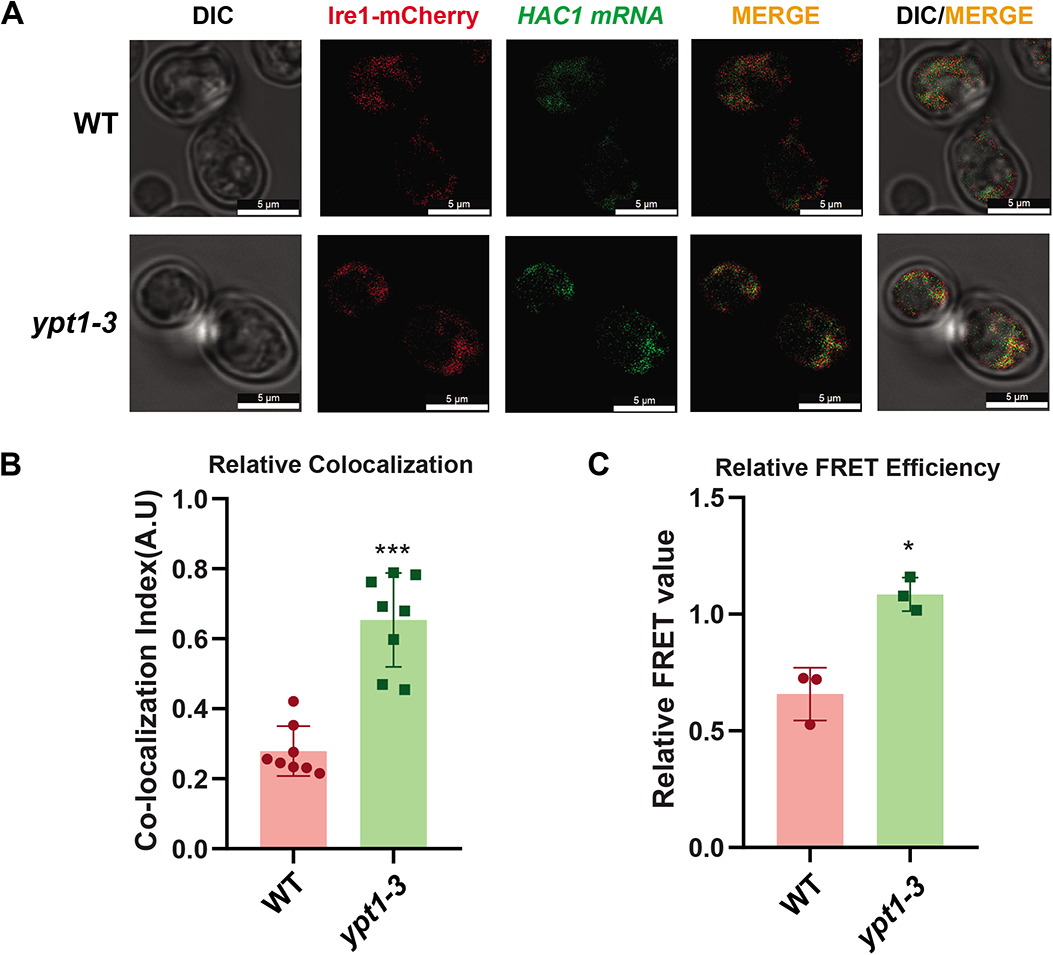
Ypt1p controls the targeting and recruitment of pre-*HAC1* mRNA to the Ire1p foci. **A.** Localization of *HAC1^NRE^* mRNA decorated with *NRE-GFP2* and Ire1-RFP following the induction of ER stress by 1 μg/ml tunicamycin in WT and *ypt1-3* yeast strains. The images were captured and processed as described in **Materials and Methods**. The bar in each field represents 5 μm. **B.** Histogram depicting the co-localization index of *HAC1^NRE^*-GFP-nucleolin mRNA signal and Ire1p-RFP signal at 10 minutes after in imposition of ER stress by 1 mg/ml tunicamycin in WT and *ypt1-3* yeast strains. Co-localization index for *HAC1^NRE^*-GFP-nucleolin recruitment into Ire1 foci was expressed in arbitrary units. Co-localization indices (Pearson correlation coefficient, PCC) were determined as described in the **Materials and Methods** section and the co-localization data is presented in supplementary table S5. **C.** Relative fluorescence resonance energy transfer (FRET) between *HAC1^NRE^*-GFP-nucleolin and Ire1p-RFP in WT and *ypt1-3* yeast strains. Quenching of donor *HAC1 ^NRE^*-GFP-nucleolin mRNA in the presence of Ire1p-RFP (acceptor) in WT and *ypt1-3* strains was determined as described in **Materials and Methods** section. Histogram depicting the Relative FRET value (in arbitrary units) from donor *HAC1^NRE^*-GFP-nucleolin mRNA into Ire1p-RFP foci at 10 minutes after ER stress by 1 µg/ml tunicamycin in a normal and *ypt1-3* strains. Donor (*HAC1^NRE^*-GFP-nucleolin mRNA) quenching as an indicator of relative FRET value was measured for *HAC1^NRE^*-GFP-nucleolin recruitment into Ire1p foci. Means and standard error of the mean were determined from n=3 (three independent biological replicates). Donor quenching estimated by Flow Cytometry from 10,000 cells of each strain were determined as described in the **Materials and Methods** section using BD FACS Verse software. The statistical significance of difference as reflected in the ranges of *p*-values estimated from Student’s two-tailed t-tests for a given pair of test strains are presented with following symbols, * <0.05, **<0.005 and ***<0.001, NS, not significant.

To bolster this finding, we estimated the extent of the co-localization of the signal intensities of *HAC1^NRE^*-Nucleolin-GFP and the Ire1p-RFP in both WT and *ypt1-3* yeast strains. The co-localization index is very low in WT yeast strain (CI = 0.28±07) (Fig. 4B, co-localization data provided in Supplementary Table S5), which is manifested by the presence of very few yellow spots (merged panel in top row in Fig. 4A). A careful examination of their distribution/localization in *ypt1-3* strain (merged panel bottom row in Fig. 4A) further revealed more intense as well as frequent co-localization of *HAC1^NRE^*-Nucleolin-GFP signal with the Ire1p-RFP (representing the cellular distribution of Ire1p foci) (CI = 0.65±0.13) (Fig. 4B, co-localization data provided in Supplementary Table S5). This finding strongly argues that a higher and more efficient targeting and recruitment of the precursor-*HAC1* mRNA to the Ire1p foci takes place in a yeast strain carrying the *ypt1-3* allele as predicted by our hypothesis. It should be noted here that the allele encoding *HAC1^NRE^*mRNA is biologically functional which was confirmed previously (32).

To further corroborate the role of Ypt1p in controlling the targeting and recruitment of pre-*HAC1* messages, we also evaluated the relative proximity of the *HAC1^NRE^*-Nucleolin-GFP module and Ire1p-YFP moiety by determining the fluorescence resonance energy transfer (FRET) from *HAC1^NRE^*-Nucleolin-GFP (donor) to Ire1p-YFP fluorophore (acceptor). Fluorescence quenching of the donor *HAC1^NRE^*-Nucleolin-GFP was determined as an indicator of FRET in WT and *ypt1-3* yeast strains at ten minutes after the addition of tunicamycin. As shown in Figure 4C, a two-fold higher donor *HAC1^NRE^*-Nucleolin-GFP quenching due to a higher FRET from *HAC1^NRE^*-Nucleolin-GFP to Ire1p-YFP was detected in the *ypt1-3* strain relative to FRET value obtained in the WT strain at ten minutes after the ER stress was imposed (Fig. 4C) (FRET efficiency data of WT and *ypt1-3* strains are provided in the supplementary Table S6). This finding, thus, strongly argues a closer physical proximity between the *HAC1^NRE^*-Nucleolin-GFP and Ire1p-YFP in *ypt1-3* strain, which in turn implies that more *HAC1* precursor mRNA is efficiently recruited onto the active Ire1p clusters in the *ypt1-3* strain. Together, these findings strongly support the conclusion that the abundance of the precursor-*HAC1* mRNA with intact BE is dictated by the Ypt1p via the reversible recruitment of the NNS/CTEXT/Exosome complex onto this precursor message followed by its subsequent decay. In WT strain, functional Ypt1p promotes the recruitment of the decay factors leading to enhanced decay resulting in a population of pre-*HAC1* transcripts most of which lack intact BE (Fig. 7). In a strain carrying a *ypt1-3* allele, the pre-*HAC1* RNA undergoes diminished exosomal degradation owing to the inability of the mutant protein to recruit NNS/CTEXT/Exosome onto this precursor RNA, which in turn facilitates the formation of a pool of precursor *HAC1* mRNAs majority of which carrying intact BE (Fig. 7). This population of pre-*HAC1* RNA with intact BE, therefore, undergoes more efficient transport from the site of transcription in the nucleus to the Ire1p foci on the ER surface. Collective evidence therefore strongly supports the view that Ypt1p dictates the targeting and recruitment of pre-*HAC1* mRNA to Ire1p foci on the ER surface during the activation of UPR, which is accomplished via the reversible recruitment of NNS-CTEXT-exosome onto the *HAC1* pre-mRNA and its subsequent decay.

### Ypt1p binds to a specific element located in the 5**′**-termini of the 3**′**-UTR of pre-*HAC1* mRNA, that facilitates its degradation by the Nuclear Decay Machinery

Having shown that, the Rab-GTPase Ypt1p plays a pivotal role in the activation dynamics of UPR signaling, we further query the molecular determinant(s) in *HAC1* pre-mRNA that triggers its susceptibility to nuclear decay factors. Previous study by Tsvetanova et al., (2012) revealed that the 3′-UTR region of the *HAC1* mRNA is necessary for Ypt1-*HAC1* interaction (33). In determining the putative Ypt1p-binding site in the 3′-UTR of the *HAC1* pre-RNA, we postulated that elimination of the specific segment(s) harboring the Ypt1p-binding site would abolish its rapid exosomal decay and prevent its participation in UPR response. Consequently, a series of deletion constructs in 3′-UTR of the *HAC1* mRNA was constructed using a combination of restriction endonuclease-digestion and reverse PCR technology (Fig. 5A). Comparison of the steady-state levels of the full-length pre-*HAC1* transcript and its various deleted versions in absence of ER-stress revealed that the levels of *HAC1-*ΔR2 and *HAC1-*ΔBE mRNA are low and similar to the physiological level of full-length message (Fig 5B). The abundance of *HAC1-*Δ3′-UTR and *HAC1-*ΔR1, in contrast, increased significantly (2 and 5 folds respectively relative to full-length mRNA) relative to the full-length message (Fig 5B). This data suggests that the region 1 (R1) spanning the 1 to 154 nt residues of the 3′-UTR of the pre-*HAC1* mRNA (Fig. 5A) plays a vital role in governing the stability of this RNA and in dictating its susceptibility to the CTEXT/nuclear exosome. Elimination of this element from the pre-*HAC1* transcript abolished its susceptibility to the nuclear exosome and CTEXT which is most likely caused by the exclusion of the putative Ypt1p binding site from this precursor mRNA. To establish whether the higher abundance of the *HAC1*-ΔR1 transcript is accompanied by its diminished decay rate, we analyzed the decay rates of full-length *HAC1* and *HAC1-*ΔR1 transcripts in the WT strain expressing functional Ypt1p. As shown in Fig. 5C, the decay rate of the *HAC1-*ΔR1 mRNA is much diminished relative to full-length *HAC1* thereby suggesting that elimination of the R1 region indeed lowered its susceptibility to the nuclear exosome/CTEXT and increased the stability of the pre-*HAC1* mRNA. Consistently, when yeast strains expressing the full-length and various deleted versions of *HAC1* pre-mRNA were challenged with tunicamycin, the strain harboring the pre-*HAC1*-ΔR1 allele exhibited better sustenance in the ER-stress more efficiently (Fig. S2C).

**Figure 5:**
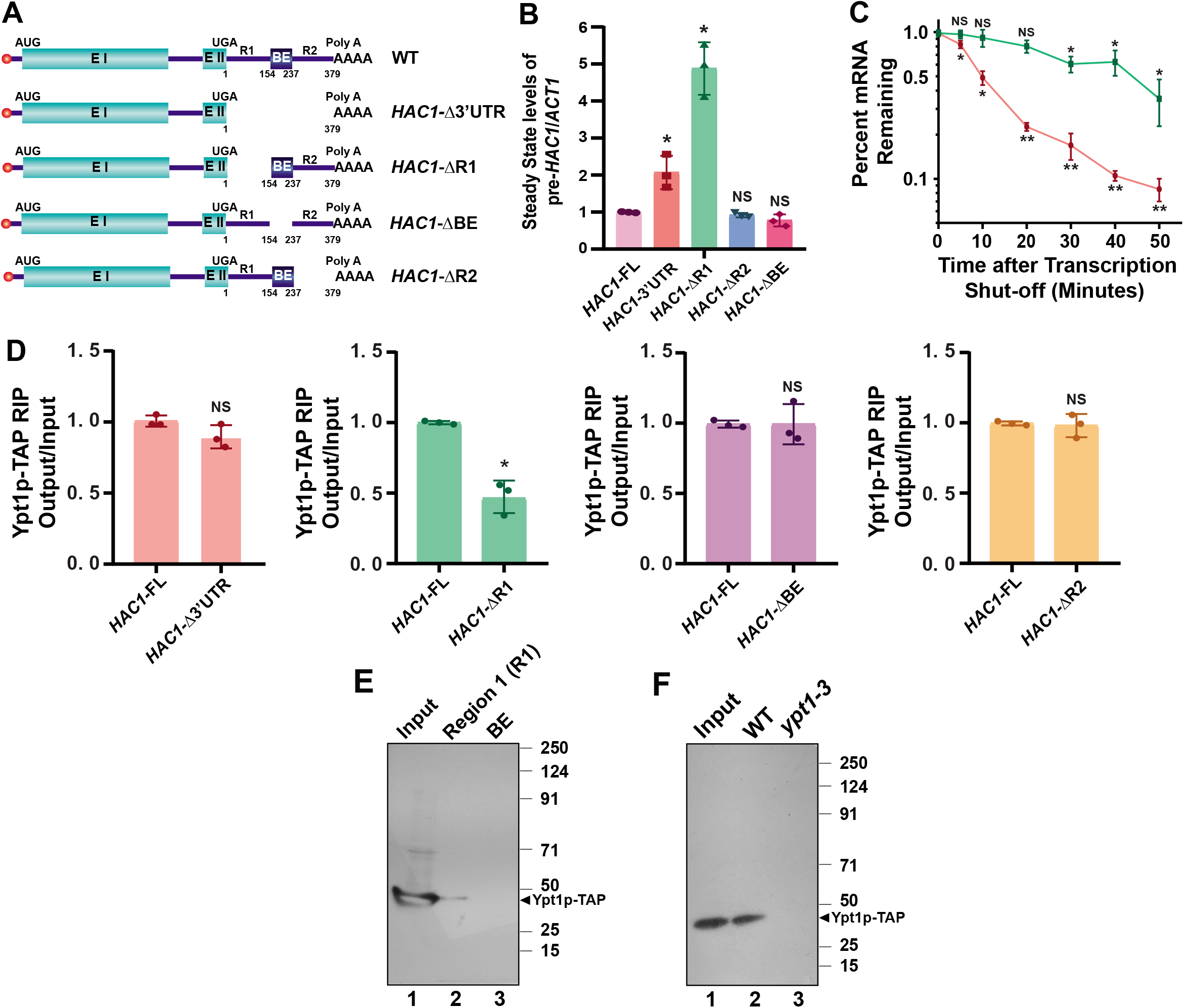
Ypt1p specifically binds to a segment encompassing first 154 nt of the 3′-UTR of pre-*HAC1* mRNA. **A.** Schematic representation of *HAC1* pre-mRNA along with its various deletions constructs used in this study. **B.** Relative steady state levels of full-length *HAC1* pre-mRNA and its various deleted versions as determined by qPCR analysis using a primer set specific to *HAC1* intron. cDNA samples were prepared from yeast strains expressing either full-length or one of its deleted version following the procedure as described in **Materials and Methods**. Transcript copy numbers/2 ng cDNA prepared from each strain was normalized to *SCR1* RNA levels obtained from respective strains and the normalized signals are presented as means ± SE (n=3 for each strain). Normalized value of full-length pre-*HAC1* mRNA was set to one. **C**. Decay rates of the full-length pre-*HAC1* (red line) and pre-*HAC1*-ΔR1 (green line) mRNAs. Decay rates were determined from yeast strains expressing either full-length or pre-*HAC1*-ΔR1 deleted version in three independent experiments (biological replicates) by qRT-PCR analysis using the primer sets spanning the *HAC1* intronic sequences. The intronic signals were normalized to *SCR1* RNA signals obtained from respective strains from each time point and normalized signals (mean values ± SD) were presented as the fraction of remaining RNA (with respect to the normalized signal at 0 min) as a function of time of incubation with transcription inhibitor 1, 10-phenanthroline. **D.** *In-vivo* binding profile of Ypt1p to the full-length and various deleted version of the pre-*HAC1* transcript. The binding was computed by qRT-PCR following immunoprecipitation by anti-TAP antibody of the whole cell extracts prepared from UV-cross-linked yeast cells expressing Ypt1p-TAP and any one of the pre-*HAC1*, pre-*HAC1*-Δ3′UTR, pre-*HAC1*-ΔR1, pre-*HAC1*-ΔBE and pre-*HAC1*-ΔR2 transcripts. Mean value from three independent experiments (biological replicates) is shown where immunoprecipitated (output) samples were normalized to input following quantification by real-time PCR. The error bars in the graph represent standard deviations (S.D.). For **B-D**, the statistical significance of difference as reflected in the ranges of p-values estimated from Student’s two-tailed t-tests for a given pair of test strains for each message are presented with following symbols, ∗<0.05, ∗∗<0.005 and ∗∗∗<0.001, NS, not significant. **E.** Ypt1p binds to the region 1 (R1) of 3′-UTR of pre-*HAC1* mRNA. Protein extracts prepared from WT yeast cells expressing Ypt1-TAP was subjected to two separate Streptotag affinity purification procedures using either *HAC1*-R1 (R1-Streptag) or *HAC1-*BE (BE-Streptotag) segment as bait. Bound pre-*HAC1*-Ypt1p-TAP RNA-protein complex was then eluted from dihydrostreptomycin coupled sepharose 6B column followed by immunobloting with anti-TAP antibody. **F**. Only the functional Ypt1p binds to the region R1 of *HAC1* pre-mRNA. Same as E except the protein extracts wer prepared either from WT cells expressing Ypt1p-TAP or from mutant strain expressing Ypt1-3p-TAP that was subjected to Streptotag affinity purification procedure using *HAC1-*R1 (R1-Streptotag) as the bait. The RNA-protein complex was eluted from dihydrostreptomycin coupled sepharose 6B column before the immunobloting with anti-TAP antibody. The position of the MW markers (in kDal) in each western blot panel is indicated at the right.

Next, we validated the functional role of the putative Ypt1p-binding element on the decay of pre-*HAC1* message by additionally investigating the effect of mutations in *YPT1* (*ypt1-3*) and *CBC1* (*cbc1-*Δ) genes on the abundance of full-length and various deleted versions of pre-*HAC1* mRNAs. As shown in Supplementary Fig S2D, the steady-state levels of *HAC1-*ΔBE and *HAC1-*ΔR2 mRNAs were enhanced in strains expressing these alleles and additionally carrying either *ypt1-3* or *cbc1*-Δ alleles relative to either isogenic WT or *HAC1*-ΔBE/ *HAC1*-ΔR2 strains (Fig S2D, lower histograms). The abundance of the *HAC1-*Δ3′-UTR and *HAC1-*ΔR1 messages, however, did not display any further enhancement in the *ypt1-3* and *cbc1*-Δ mutant yeast strains (Fig S2D, upper histograms). This finding strongly indicates that both *HAC1-*ΔR1 and *HAC1-*Δ3′-UTR mRNAs lacking the Ypt1p binding element failed to recruit the nuclear decay machinery in absence of ER stress leading to their diminished degradation and thus mutations in the *YPT1* gene did not have any further effect on their stability (supplementary Fig. S2D). In contrast, *HAC1-*ΔBE and *HAC1-*ΔR2 messages still harboring the Ypt1p-binding site underwent a rapid Ypt1p-dependent degradation in the nucleus in absence of ER stress and displayed an increase in their steady-state levels in *ypt1-3* and *cbc1*-Δ mutant backgrounds. Collective data thus strongly indicate the Ypt1p-binding element appears to be present within the first 154 bp region of *HAC1* 3′-UTR, which is responsible for its exosome/CTEXT-dependent nuclear decay.

Next, we validated the binding of the Ypt1p to the full-length and diverse deleted versions of the pre-*HAC1* mRNAs *in vivo* using RNA immunoprecipitation (RIP). As performed before, RIP signals were estimated from the extract prepared from yeast strains expressing Ypt1p-TAP and either a full length (FL) or a deleted version of the pre-*HAC1* mRNA (Fig. 5D). As shown in Fig. 5D, a significantly lower RIP signal was noted in *HAC1-*ΔR1 transcripts lacking the Ypt1p binding-site relative to full-length (FL) RNA, whereas a similar and comparable RIP-signal were observed for both of the *HAC1-*ΔR2 and *HAC1-*ΔBE mRNAs, affirming that these mutant versions still harbor the Ypt1p binding site.

To strengthen the above finding, we subjected the R1 segment of the pre-*HAC1* 3′-UTR harboring the Ypt1p-binding element to a Streptotag-based affinity purification system (38) to inquire if the Ypt1p binds this RNA element *in vitro*. For this experiment, two different sequences corresponding to regions R1 and BE were separately fused to a Streptotag aptamer by two independent PCR reactions (Supplementary Fig. S3). Each of these reactions consisted of a sense primer consisting of a T7 promoter fused in tandem to 5′-segment of either R1 or BE regions and an anti-sense primer consisting of the 3′-segments of either R1 or BE region fused in tandem to Streptotag aptamer sequence using a cloned version of *HAC1* gene as a template (Supplementary Fig. S3). The amplified products from each of these reactions were further used as the templates for two separate *in vitro* transcription reactions using T7 RNA polymerase and the fusion RNAs thus yielded (Supplementary Fig. S3) were subjected to two independent coupling reactions to epoxy-activated Sepharose 6B resin in presence of 1 mM dihydrostreptomycin (Supplementary Fig. S4B) One mg of total protein extract prepared from yeast strain expressing Ypt1p-TAP were mixed separately and independently with the two immobilized RNA-streptotag (either R1-streptotag or BE-streptotag) aptamer complexes in sepharose 6B column (Supplementary Fig. S4B). The columns were washed with tRNA and finally, the bound protein complexes were eluted with streptomycin (Supplementary Fig. S4B). The input and the two eluate samples from each column were subjected to western blot analysis using an anti-TAP antibody (Ypt1p TAP-tagged). As shown in Fig. 5E, Ypt1p was detected in both the input and 1^st^ eluate samples containing the 154 nt sequence corresponding to the R1 harboring the Ypt1p binding site as the bait. No Ypt1p was detected in the 2^nd^ eluate sample that contained the 83 nt BE region as the bait (Fig. 5E). Furthermore, in another independent experiment, separate protein extracts prepared from yeast strains expressing either functional Ypt1p or mutant Ypt1-3p were also subjected to Streptotag affinity purification with R1-streptotag as a bait. As shown in Fig. 5F, no signal was detected in the extract prepared from the strain expressing the mutant protein (Fig. 5F). Data from these experiments strongly confirmed that Ypt1p binds to the 154 nt *cis*-acting element corresponding to the region 1 (R1) of the 3′-UTR of pre-*HAC1* mRNA both *in vitro* and *in vivo* and the binding is very specific. Collective findings thus revealed that in the absence of ER-stress, Ypt1p binds to the pre-*HAC1* message in a region spanning the residues 1-154 (assuming the TGA of the *HAC1* ORF as 0 and the very next nucleotide in the 3′-UTR as +1) of its 3′-UTR that in turn further recruits NNS, CTEXT and exosome complex and thereby promoting the rapid decay of this precursor RNA.

### A large fraction of cellular Ypt1p localizes to the nucleus in the absence of ER stress, which redistributes to the cytoplasm when ER stress is imposed

Next, we investigated the cellular distribution of Rab-GTPase Ypt1p in both ER stressed and unstressed conditions to determine whether Ypt1p localizes to the nucleus. The protein was expressed from its native promoter as C-terminally TAP-tagged fusion and its intracellular localization was observed by indirect immunofluorescence followed by confocal imaging. As shown in Fig. 6A, in the unstressed condition, a large fraction of Ypt1p was indeed found to localize into the nucleus and the degree of its nuclear localization is significantly higher relative to its cytoplasmic distribution (Fig.6A). In contrast, in ER-stressed condition, Ypt1p displayed more dispersed cytoplasmic distribution with dramatically less localization in the nucleus in all visible fields. Analysis of the co-localization index between the Hoechst (nucleus) and the Ypt1p-FITC signals revealed a significantly (2 folds) higher co-localization index during the unstressed condition that was drastically reduced when ER-stress was imposed (Fig. 6C) (Co-localization data of this experiment is provided in Supplementary Table S7). Interestingly, the mutant Ypt1-3p protein did not display a significant amount of nuclear localization either in stressed or unstressed conditions (Fig. 6B). Insignificant amount of nuclear localization of the mutant Ypt1-3p was noted during the unstressed condition (Fig. 6B, upper panel) which was consistently reflected in its co-localization index during both conditions (Fig. 6D) (Co-localization data of this experiment is provided in Supplementary Table S7). This interesting finding, therefore, suggests that in absence of ER-stress, Ypt1p primarily localizes in the nucleus whereas ER stress promotes its rapid redistribution to cytoplasm leading to a significant exclusion from the nuclear territory (Fig. 6A, Merge Panel). The majority of the mutant Ypt1-3p was unable to localize to the nucleus and failed to bind to the pre-*HAC1* message, which in turn led to its failure to promote the nuclear decay of this precursor RNA. Preferential and reversible nuclear localization of Ypt1p during the unstressed condition, thus, provides a mechanism by which Ypt1p-association with pre-*HAC1* mRNA may be regulated during UPR signaling dynamics in baker’s yeast.

**Figure 6:**
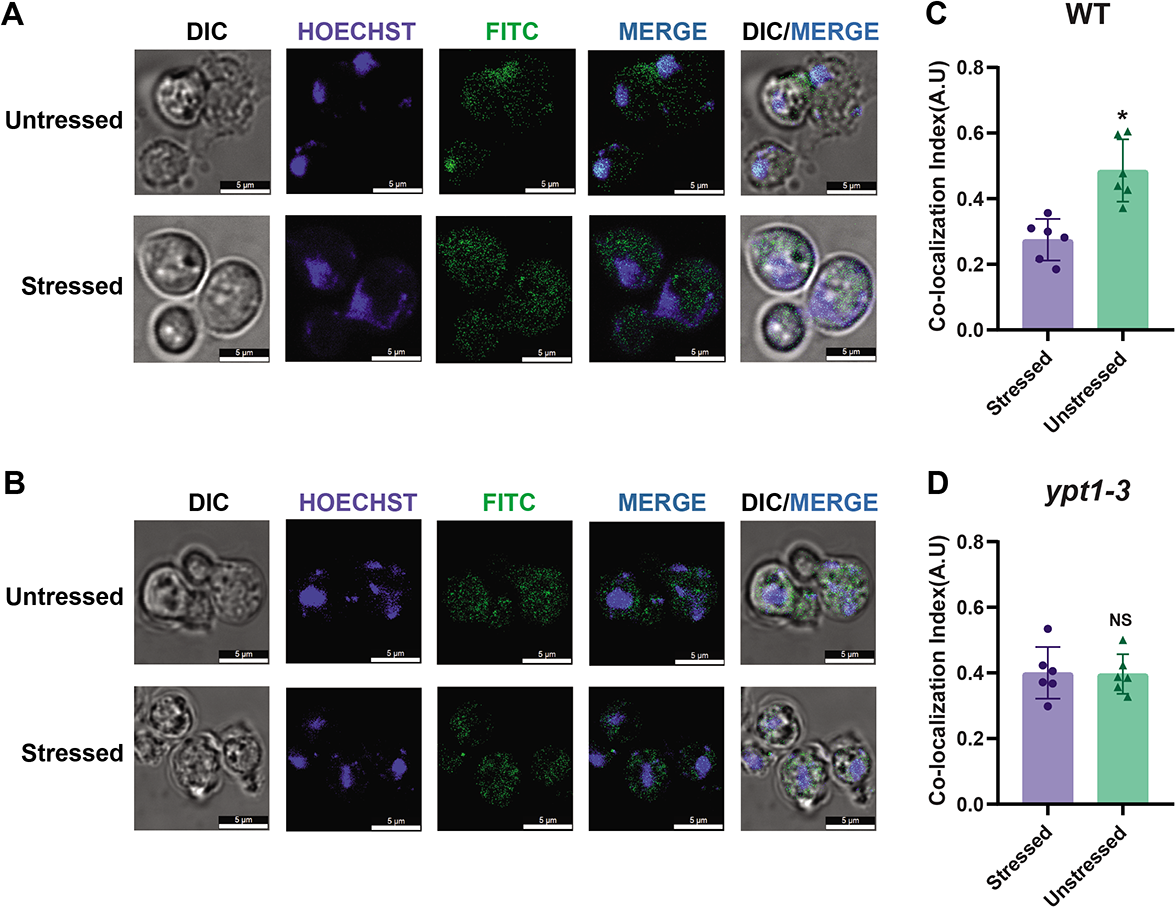
Ypt1p relocalizes to the cytoplasm during the ER stress. **A-B.** Localization of C-terminally TAP tagged WT Ypt1p **(A)** and mutant Ypt1-3p **(B)** expressed from their native promoters as determined by immunofluorescence using rabbit anti-TAP primary antibody followed by anti-rabbit FITC conjugated secondary antibody in both unstressed (upper panel) and ER-stressed (induced by 1µg/ml Tunicamycin) (lower panel) condition. The nucleus of both samples were stained with Hoechst stain. The bar in each field represents 5 μm. **C-D.** Histogram depicting the co-localization of either WT Ypt1p FITC signal/Hoechst (nucleus) **(C)** or mutant Ytp1-3p FITC signal/Hoechst (nucleus) **(D)** before and after ER stress was induced. Co-localization indices (Pearson correlation coefficient, PCC) were expressed in arbitrary units and were determined as described in the **Materials and Method** section. Co-localization data for this figure is presented in supplementary Table S5. The statistical significance of difference as reflected in the ranges of *p*-values estimated from Student’s two-tailed t-tests for a given pair of test strains for different conditions are presented with following symbols, * <0.05, **<0.005 and ***<0.001, NS, not significant.

## DISCUSSION

Diverse extracellular insults and aberrations in protein folding result in a disparity in the normal protein folding activity of ER thereby leading to the accumulation of misfolded proteins in ER lumen, dubbed “ER stress”. The unfolded protein response (UPR) readjusts ER folding capacity to restore protein homeostasis within ER lumen. The key transcription activator, Hac1p responds to ER stress by activating the genes encoding ER chaperones, which collectively refold the unfolded ER proteins and thereby restore ER homeostasis. Previous investigation from our lab revealed that the efficiency of the delivery and recruitment of the precursor *HAC1* to the Ire1p foci for its successful non-spliceosomal splicing is governed by the reversible and dynamic nuclear decay of this precursor message that is dependent on the nuclear exosome and CTEXT (32). Preferential nuclear mRNA degradation of this precursor RNA thus plays a pivotal role in tuning the gain of the UPR signaling dynamics (32). Nevertheless, the precise mechanism of specific and selective recognition of *HAC1* pre-mRNA from the vast majority of other nuclear messages by the nuclear decay machinery remained obscure. Recent evidence of association of the Rab-GTPase Ypt1p to unspliced *HAC1* message that leads to alteration of its stability (33) inspired us to explore if Ypt1p plays any key role in the recognition and targeting of the *HAC1* pre-mRNA.

Our data demonstrated that indeed Ypt1p plays an important role in controlling the UPR signalling dynamics in *Saccharomyces cerevisiae*, which validated the earlier findings reported by Tsvetanova et al. (2012) (33). While these authors employed an *in vitro* proteomic analysis approach to identify the physical association between *HAC1* and Ypt1p, we utilized a genetic approach using a specific mutant allele *ypt1-3* to investigate its role in UPR response. Our initial finding revealed that the yeast strain carrying the *ypt1-3* mutant allele displayed better sustenance to ER stress when challenged with tunicamycin and DTT (Fig. 1A-C). Further investigation showed that Ypt1p impacts the steady-state levels of precursor *HAC1* mRNA by altering its stability both in the absence and presence of ER stress (Fig. 1). Notably both the growth and pre-*HAC1* stability phenotypes displayed by *ypt1-3* mutant cells are identical to those shown by the mutants deficient in the nuclear mRNA decay thereby strongly indicated that Ypt1p participates in the same regulatory pathway that controls the rapid nuclear decay of Pre-*HAC1* RNA. Subsequent validation of this postulate by analyzing the decay rate of the pre-*HAC1* message established that Ypt1p protein indeed facilitates the accelerated decay of this precursor transcript (Fig. 1E-F).

This finding prompted us to address if Ypt1p functions in the initial recognition of pre-*HAC1* mRNA by promoting the selective and sequential recruitment of the exosome-specificity factor NNS complex followed by the nuclear decay factors onto pre-*HAC1* mRNA. As shown in Figs 2-3, Ypt1p displayed a strong association with the representative components of NNS, exosome, and CTEXT complex in the absence of stress, which was reversed when ER stress was imposed. Subsequent analysis to detect if Ypt1p facilitates the accelerated decay of pre-*HAC1* message by coordinating the sequential selective recruitment of NNS, exosome, and CTEXT components onto this RNA during its transcription indeed validated our prediction. Our collective data strongly argues in favour of the view that indeed Ypt1p promotes the co-transcriptional recruitment of the NNS complex onto the precursor *HAC1* transcript, which in turn, further recruits the CTEXT and the nuclear exosome sequentially onto it to trigger its preferential nuclear decay (Fig. 7). Recruitment of Ypt1p onto pre-*HAC1* mRNA in a preferential manner thus provides a mechanism by which this precursor message is initially recognized by the decay machinery in a selective fashion from the vast majority of normal and other aberrant messages. Physical association of Rab-GTPase Ypt1p with pre-*HAC1* mRNA, therefore, provides a means by which the initial recognition of this pre-mRNA may be accomplished.

**Figure 7:**
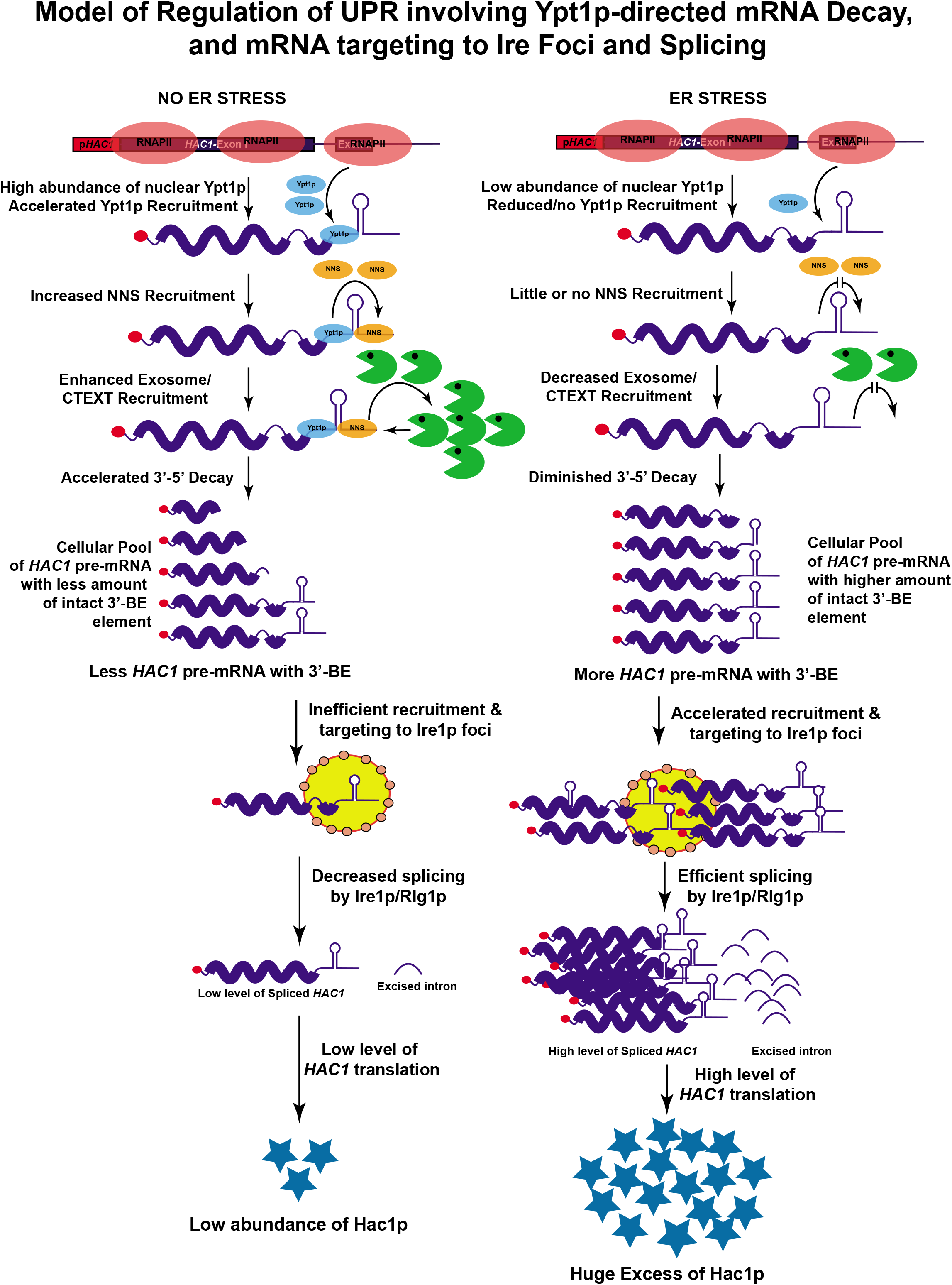
Model of Ypt1p-directed regulation of UPR in baker’s yeast. Left Side. In absence of ER-stress, nuclear Ypt1p is co-transcriptionally recruited on to precursor *HAC1* message in a selective fashion that further promotes the co-transcriptional and sequential recruitment of NNS, CTEXT and the nuclear exosome onto this this precursor message and trigger its dynamic and reversible 3′→5′ degradation. This rapid decay results in the formation of a *HAC1* precursor RNA pool most of which lack the intact BE leading to their less efficient delivery to Ire1p foci, less splicing and translation. **Right Side.** ER-stress triggers the relocalization of Ypt1p from nucleus to cytoplasm leading to its inefficient recruitment onto the precursor *HAC1*, which in turn affects the loading of the decay factors on this precursor RNA resulting in its diminished decay. Diminished pre-*HAC1* decay leads to the formation of a pool of *HAC1* pre-RNA with an intact and functional BE, which are targeted and recruited more efficiently to the Ire1p foci leading to more efficient splicing and translation.

Consistent with the characteristic directionality of the exosomal degradation of pre-*HAC1* mRNA in the 3′→5′ direction that typically initiates from the extreme 3′-termini of the precursor message, our data suggests that Ypt1p binds to the region 1 (R1) located at the extreme 5′-terminus (encompassing 1-154 nt relative to TGA codon) of the 3′-UTR of pre-*HAC1* mRNA. Interestingly, Ypt1p binds to the R1 region both *in vivo* (using RIP technique) and *in vitro* (using Streptotag affinity purification system) (Fig. 5D-F). Importantly, neither the mutant Ypt1-3p protein displayed any binding to the R1 region nor did the functional protein showed any affinity to BE region in our Streptotag procedure. Moreover, the stability of one of the deleted versions of *HAC1* pre-RNA, the *HAC1*-ΔR1 displayed much higher stability relative to the full-length *HAC1* precursor in the WT yeast strain equipped with functional NNS complex and nuclear decay machinery (Fig. 5B-C). Collective data therefore strongly affirmed that the binding of Ypt1p to the region R1 is extremely specific and physiologically relevant. Once Ypt1p binds to the region R1 in the absence of ER-stress, it further recruits the NNS complex presumably in the vicinity of the *HAC1* pre-mRNA, which in turn assists loading of the nuclear exosome and CTEXT presumably at the extreme 3′-termini for its dynamic and reversible decay (Fig. 7).

In good agreement with the nuclear role of Ypt1p, our data suggests that it is a shuttling protein. In the absence of ER-stress, a majority of cellular Ypt1p is distributed in the nucleus and upon imposition of ER-stress, it redistributes itself to the cytosol (Fig. 6). Although, the exact mechanism of reversible binding of Ypt1p to the pre-*HAC1* mRNA during various phases of ER-stress is currently unknown, our finding of the relocalization of the Ypt1p to the cytosol during the activation phase of the UPR suggests that at least this reversibility could partly be accomplished by its relocalization from the nucleus to the cytoplasm. However, it is currently unclear what factor drives Ypt1p from the nucleus to the cytoplasm when ER-stress is enforced. In addition, other factors may also be involved in altering the affinity of the Ypt1p in binding the pre-*HAC1* mRNA in presence of ER stress. Future research should unfold the mechanism of Ypt1p redistribution during various phases of UPR, such as activation and attenuation.

Our finding of the existence of a separate layer of regulation in the control of cellular UPR signalling dynamics suggests a possible mechanism by which a small extent of signalling cue can amplify the final output of this pathway optimally. It is noteworthy that protein-unfolding status within the ER lumen may be sensed by two different sensors, which are eventually transduced into the massive production of Hac1p (Fig. 8). While one sensor of protein unfolding is Ire1p, the other sensor is probably Ypt1p. As reported by numerous laboratories, the sensing of the unfolding status by Ire1p is followed by its oligomerization leading to the activation of its RNase domain that eventually splices out the *HAC1* intron (28, 29, 40–42). Sensing by Ypt1p, in contrast, is manifested by its quick relocalization from the nucleus to the cytoplasm thereby causing its decreased recruitment onto pre-*HAC1* mRNA resulting in its diminished decay and increased delivery to Ire1p foci (Fig. 8). The discovery of this second regulatory circuit involving nuclear mRNA decay that functions upstream of the Ire1p sensing circuit is a unique characteristic of this signalling mechanism. The existence of the Ypt1p-exosome circuit indicates that during the activation of UPR, the outputs of two independent circuits converge on the Ire1p interface on ER surface thereby magnifying the final transduced output in a synergistic fashion (Fig. 8). Ypt1p thus plays a vital role in sensing the ER-stress signal independently and transduce that signal to the Ire1p kinase-endoribonuclease via the increased delivery of pre-*HAC1* mRNA during the activation phase of ER-stress cycle.

**Figure 8:**
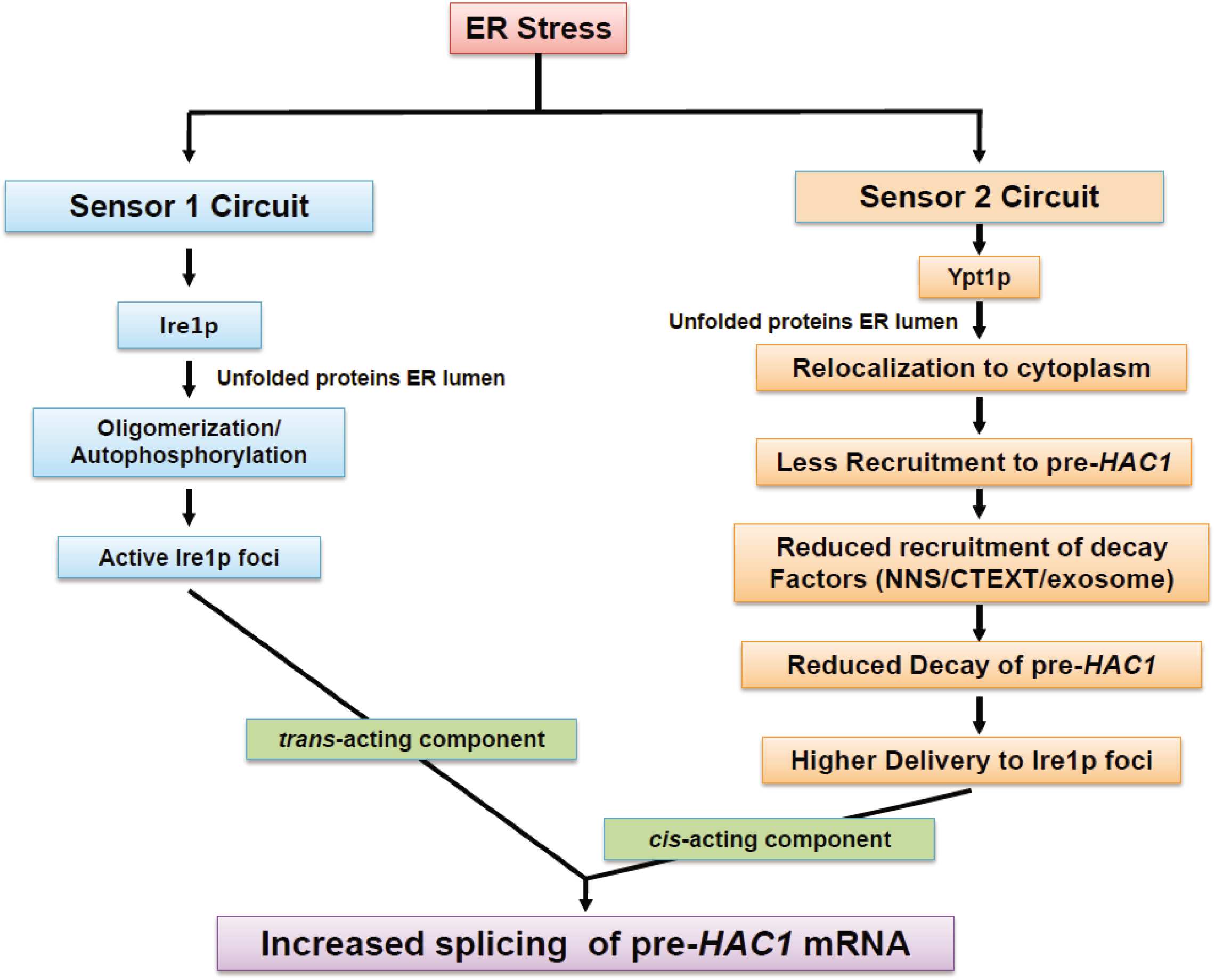
Two independent regulatory circuits control of activation of UPR signaling dynamics in baker’s yeast. Cartoon depicting two different sensors that sense the unfolded proteins within the ER lumen during the activation stage of UPR. While Ire1p serves as one of the sensors, Ypt1p plays a crucial role as the other sensor. Sensing by Ire1p is characterized by its oligomerization and formation of an active RNase domain that eventually splice out the *HAC1* intron. Sensing by Ypt1p, in contrast, is accompanied by its quick relocalization from the nucleus to the cytoplasm thereby causing its decreased recruitment onto pre-*HAC1* mRNA resulting in its diminished decay and increased delivery to Ire1p foci. The timely convergence of the output signals from two independent circuits on the Ire1p interface on ER surface during the activation phase of ER-stress results an amplified an integrated signal that eventually produces copious amounts of functional *HAC1* mRNA and Hac1p protein.

## EXPERIMENTAL PROCEDURES

### Nomenclature, strains, media, and yeast genetics

Standard genetic nomenclature is used to designate wild-type alleles (e.g., *YPT1*, *HAC1*, *NRD1, CYC1*, *RRP6*), recessive mutant alleles (e.g., *ypt1-3*, *nrd1-1,*) and disruptants or deletions (e.g., *cbc1*::*TRP1*, *cbc1-*Δ, *rrp6*::*TRP1*, *rrp6*-Δ). An example of denoting a protein encoded by a gene is as follows: Ypt1p encoded by *YPT1*. The genotypes of *S. cerevisiae* strains used in this study are listed in **Table S1**. Standard YPD, YPG, SC-Lys (lysine omission), and other omission media were used for testing and growth of yeast (43). Yeast genetic analysis was carried out by standard procedures (43).

### Plasmids and oligonucleotides

The plasmids were either procured from elsewhere or were constructed in this laboratory using standard procedures. All the oligonucleotides were obtained commercially from Integrated DNA Technology (Coralville, IA, USA). The plasmids and oligonucleotides used in this study are listed in **Tables S2 and S3**, respectively.

#### Construction of TAP tagged strains

The yeast strains harbouring the TAP-tagged version of WT Ypt1p and mutant Ypt1-3p proteins were generated by amplifying the TAP tag sequence along with the flanking region of the Ypt1p protein using the pBS1539 plasmid as the template (44). The PCR product was integrated in the genomic Ypt1p or Ypt1-3p loci following the transformation into appropriate yeast strains using two-step gene replacement.

#### Construction of various deleted versions *HAC1* allele

For the construction of various deleted versions of 3′-UTR of *HAC1* allele, specific primers flanking the region of interest are oriented facing away from each other such that the entire fragment of DNA excepting the specific region gets amiplified. A *CEN* plasmid harbouring a full length *HAC1* is used as template and Q5® High-Fidelity DNA Polymerase (New England Biolabs Inc., MA, USA) is used. The reaction is set according to manufacturer’s protocol. The amplicon thus obtained is purified from gel and ligated using T4 DNA Ligase (New England Biolabs Inc., MA, USA) and maintained in *HAC1-*Δ background.

#### Cell Viability Assay

Cell Viability Assay was performed as described previously (45). Briefly, each strain was grown overnight, and the culture was then subsequently diluted to 1 × 10^7^ cells/ml. This stock was further diluted 10 fold serially to achieve 1 × 10^4^, 1 × 10^3^, and 1 × 10^2^ cells/ml respectively. 3 µl of each stock was then spotted onto the surface of YPD plate (Yeast Extract 1%, Peptone 2%, Dextrose 2%, Agar 2%) without and with 1µg/mL tunicamycin (Sigma-Aldrich Inc., MI, USA). Plates were then incubated for 48-72 hours at 30°C before capturing their photographs. For liquid growth assay, specific strains of *S. cerevisiae* were grown in triplicate until the O.D._600_ reaches 0.6 when either DMSO (mock treated, -Tm/-DTT) or 1µg/mL tunicamycin (+Tm) or 5mM DTT (+DTT) (Thermo Fisher Scientific Inc., MA, USA) was added to one of them. Aliquots of each culture were collected at different times after addition of tunicamycin or DTT, appropriately diluted and subsequently spreaded onto YPD plates. Plates were then incubated for 48-72 hours at 30°C and the total number of colonies for a specific strain were counted and plotted as histogram.

#### RNA Analyses and Determination of Steady-State and decay rate of mRNAs

Total RNA was isolated as described earlier (18) by harvesting appropriate yeast strains followed by extracting the cell suspension in the presence of phenol-chloroform-IAA (25:24:1) and glass bead. Following the extraction, the RNA was recovered by precipitation with RNAase-free Ethanol.

For the preparation of cDNA, total RNA samples were first treated with 1 µg RNAase free DNase I (Fermentas Inc., MA, USA) at 37°C for 30 minutes followed by first strand cDNA synthesis using Superscript Reverse Transcriptase (Thermo Fisher Scientific Inc., MA, USA) using Random Primer (Bioline Inc.) by incubating the reaction mixture at 50°C for 30 minutes. Real Time qPCR analyses with 2 to 3 ng of cDNA samples were used to determine the steady-state levels of pre-*HAC1*, *HAC1,* and *SCR1,* etc. were performed as described previously.

Stabilities of a specific mRNA was determined by the inhibition of global transcription with transcription inhibitor 1, 10 phenanthroline (Sigma-Aldrich Inc., MI, USA) at 30°C, as described (18) previously. Briefly, the specific strain was grown at 30°C till mid-logarithmic phase, 1,10 phenanthroline was then added to the growing culture at 20 µg/ml final concentration followed by withdrawal of a 25 ml of aliquot of culture at various times after transcription shut off. Messenger RNA levels were quantified from cDNA by Real-time PCR analysis and the signals associated with the specific message was normalized against *SCR1* signals. The decay rates and half-lives of specific mRNAs were estimated with the regression analysis program (Graphpad Prism version 8.0.1) using a single exponential decay formula (assuming mRNA decay follows a first order kinetics), y =100 e^-bx^ was used.

#### Immuno Blot Analysis

Total protein was isolated from specific yeast strains grown overnight at 30°C in YPD broth. Following the harvesting by centrifugation at 5000 rpm for 7 minutes the cell pellets were quickly frozen in liquid nitrogen and stored at −70°C. Frozen pellets were thawed on ice and resuspended in 1 ml of Buffer A (50 mM Tris-Cl pH 7.5, 150 mM NaCl, 5 mM EDTA, 1 mM DTT, 1 mM PMSF) supplemented with Protease inhibitor (Invitrogen Inc. Carlsbad, CA, USA) and the cells were lysed by vortexing 10-15 times with glass beads followed by clarification of the particulate fraction by centrifugation. Supernatants were collected by centrifugation at 10, 000 rpm for 20 minutes and saved as the total soluble protein fraction for further analysis. Protein concentration was determined by Bradford reagent assay kit (Bio-Rad Inc., Valencia, CA, USA). 30-50 µg of total protein was used, which was resolved either in a 8% or a 10% SDS-polyacrylamide gel and then immunoblot analysis using primary antibodies for specific proteins for 1 hr at room temperature in following dilutions: rabbit polyconal anti-TAP (cat.#CAB1001) (1:1000), rabbit polyclonal anti-Cbc1 (1:1000), rabbit polyclonal anti-Rrp6 (1:1000), rabbit polyclonal anti-Tif4631(1:2000), rabbit polyclonal anti-Rrp4 (1:1000). Immuno-reactive bands were developed and detected by chemiluminescence (ECL imager kit, Abcam) and the images were captured either by X-ray film or myECL Chemidoc Imager (Thermo Scientific, USA).

#### Chromatin Immunoprecipitation (ChIP)

Chromatin preparation was performed as described earlier (46). The *YPT1* and *ypt1-3* strains or *NRD1* and *nrd1-1* strains in different background were used for this study. 150 ml of cells grown until OD_600_ ≈ 0.5 (≈10^7cells/ml) and were fixed with 1% formaldehyde for 20min. After glycine addition to stop the reaction, the cells were washed and lysed with glass beads to isolate chromatin. The cross-linked chromatin was sheared by sonication to reduce average fragment size to≈500bp. Immunoprecipitation of chromatin was carried using ChIP assay kit (EZ-ChIPTM; 17-295, MilliporeSigma, MA, USA). Immunoprecipitation of 120 μg Chromatin fractions (≈100μl) from each strain was performed with protein G agarose and specific Antibody overnight at 4◦C. After washing beads and chromatin elution, the eluted supernatants and the input controls were incubated with Proteinase K for 1h at 37◦C followed by 5h at 65°C to reverse cross-linked DNA complexes. DNA was extracted using the DNA-clean up column provided with the kit. The immunoprecipitated DNAs (output) were quantitated by real time PCR using specific primers located along a specific gene coding sequence and normalized to a half dilution of input DNA. Amplifications were done in duplicate for each sample, averages and standard deviations were calculated based on three independent experiments.

#### Analysis of Binding Profile of mRNA by RNA Immunoprecipitation

Yeast strains containing TAP-tagged *YPT1* cells or other cells were grown to O.D._600_ 0.6 to 0.8. Prior to the lysis of the cells intact yeast cells were cross-linked by UV-irradiation in a petri dish on ice. Cells were washed once with Buffer A (50 mM Tris-HCl pH 7.4, 140 mM NaCl, 1.8 mM MgCl_2_, 0.1% NP-40) then resuspended in Buffer B (Buffer A supplemented with 0.5 mM DTT, 40 U/ml RNase Inhibitor, 1 mM PMSF, 0.2 mg/mL heparin, protease inhibitor) and lysed by vortexing using glass beads. Lysates were cleared by centrifugation for 10 min at 8,000 rpm/4°C. The lysate was quantified and pre-cleared for 30 minutes at 4°C. Protein A sepharose beads (Santa Cruz Biotechnology, TX, USA) were coated for overnight at 4°C using TAP antibody (Cat#CAB1001) (Thermo Fisher Scientific, MA, USA) in Buffer B. One mg pre-cleared cell lysate was incubated with antibody-coated beads for 4 hours at 4°C on a rotator. Beads were washed twice in Buffer B for 5 min/4°C/rotator each, and once in Buffer C (50 mM Tris pH-7.4, 140 mM NaCl, 1.8 mM MgCl_2_, 0.01% NP-40, 10% glycerol, 1 mM PMSF, 0.5 mM DTT, 40 U/ml RNase Inhibitor, protease inhibitor) for 5 min/4°C/rotator each. After washing, beads were resuspended in 200 μl Buffer C supplemented with 1% final SDS and heated at 70°C for 45 min with constant mixing to de-crosslink samples and elute antibody-bound protein-RNA complexes from the beads. Cells for ‘‘Mock IP’’ from isogenic parental untagged strain were grown, cross-linked, lysed, and incubated with anti-TAP antibody and protein A plus sepharose beads following the same procedure. Finally RNA was isolated from the eluates by phenol-chloroform-Isoamyl alcohol (25:24:1) pH 4.5 extraction and cDNA was prepared using 1^st^ strand cDNA synthesis kit (Takara Bio Inc., Shiga, Japan) as described in above RNA analysis section. The immunoprecipitated RNAs (output) were quantitated by real time PCR using specific primers located along a specific gene coding sequence and normalized to a half dilution of input DNA. Amplifications were done in duplicate for each sample, averages and standard deviations were calculated based on three independent experiments.

#### Co-Immunoprecipitation

Cells were grown in 50ml YPD until OD_600_ 2.7-3.0 is reached. Cell lysate was prepared as described previously in 1ml of Buffer A. Protein A^PLUS^ Agarose Beads (Santa Cruz Biotechnology, TX, USA) ∼20 μl bed volume per reaction were equilibrated twice with ten volumes of Buffer A. The beads for each washing was resuspended in Buffer A and kept on a rotator wheel for 5 mins at 4°C. The beads were centrifuged each time to remove residual Buffer A. The extract was then pre-cleared by adding the equilibrated beads and incubation on a rotator wheel at 4°C for 30 minutes followed by centrifugation to get rid of the beads. To the pre-cleared protein extract specific antibody was added and incubated for 4 hours on rotator wheel at 4°C. To this extract, 50 μl bed volume of pre-equilibrated Protein A^PLUS^ Agarose Beads was added and incubated at 4°C on rotator wheel overnight. The beads are washed thrice with ten volumes of Buffer A by rotating on rotator wheel at 4°C for ten minutes each. Finally, elution was performed by boiling the beads in 40 μl of SDS loading dye for 5 minutes. Samples were then analyzed on 12% SDS-PAGE gel followed by Western blot analysis.

#### Localization of Ypt1p by indirect immunofluorescence

Cells were grown in 10 ml culture tubes up to mid-log phase followed by fixing them in 4% paraformaldehyde solution for 30 minutes after getting rid of the residual culture media. Cells are then washed twice in 0.1 M KPO_4_ buffer and once in 0.1M KPO_4_/1.2M Sorbitol and finally resuspended in 1ml 0.1 M KPO_4_/1.2M Sorbitol. The cells were then spheroplasted for permeabilization by mixing 0.5 ml cells to 5 μl of 10mg/ml Lyticase (Sigma-Aldrich Inc., MI, USA) and 10 μl β-mercaptoethanol (Sigma-Aldrich Inc., MI, USA) followed by incubation at 30°C for 10-20 minutes as per cell density. The reaction was supplemented with 10 μl of β-mercaptoethanol for optimal reaction. After adequate amount of spheroplasting, cells are centrifuged at 5000 rpm for 1 minute and washed gently with 0.1 M KPO_4_/1.2M Sorbitol twice and finally resuspended in 200 μl of KPO_4_/1.2M medium. Cells are now ready to be adhered onto poly-lysine coated cover-slips (regular glass-slips are appropriately poly-lysine coated according to Manufacturer’s protocol). About 100 μl of spheroplasted cells are added onto the poly-lysine coated cover-slips and allowed to sit for 15-20 minutes. After aspirating out the supernatant cells are blocked with 400 μl of 10% PBS-BSA for 30 minutes followed by the addition of primary antibody (anti-TAP antibody at 1:200). Cells are incubated overnight at 4°C in humid chamber. The cells were then washed five times with PBS-BSA before incubation with secondary antibody (Goat anti-rabbit secondary antibody, FITC, Cat #F-2765 at 1:1000) at room-temperature in the dark for 1hour. The cells were washed further four times with PBS-BSA followed by incubating the cells with 1.5 µg/ml Hoechst to stain the nucleus. The cells were then washed extensively with PBS-BSA and finally mounted onto a clean glass slide using 10 mg/ml p-phenylenediamine (Sigma) in 90% glycerol. The cells were observed in Confocal Laser Scanning Microscope (Leica) and the localization of *YPT1*was analyzed from the FITC.

#### Localization analysis of *HAC1^NRE^-GFP* mRNA and Ire1p-RFP proteins using confocal microscopy and Image Processing

The co-localization imaging of *HAC1* mRNA with Ire1 proteins in the live cell was done by using three plasmids (37) as described earlier (32). The reporter plasmid carries an engineered version of *HAC1* mRNA in which a 22-nucleotide RNA module, identical to the nucleolin recognition element (NRE) was inserted at the 3’-UTR (*HAC1^NRE^*). The second plasmid expressed a chimeric protein consisting of an NRE-binding nucleolin domain (ND) which is fused to the monomeric form of the green fluorescent protein 2 (GFP2). In the third plasmid, Ire1p is conjugated with red fluorescent protein (RFP). All these plasmids were transformed into appropriate yeast strain and was grown until the OD_600_ reaches 0.6. 1 µg/mL of Tunicamycin was then added to the medium and incubated for another 1 hour at 30°C. An aliquot of the cell was used for *in vivo* imaging studies. Images were captured using Confocal Laser Scanning Microscope (Leica). The GFP and RFP were excited at 458 nm and 514 nm respectively, and emissions were recorded at 466-526 nm and 530-650 nm respectively. Image processing and analysis of co-localization Index were done using Leica Microsystem software.

#### Flow Cytometry-Based FRET

Flow Cytometry-based Fluorescence Resonance Energy Transfer (FRET) measurement was done by using the protocol as described earlier (27, 42). The amount of signal quenched in the presence of acceptor from a donor is considered as the indicator of quantitative FRET and donor-acceptor pair should have significant overlap of excitation spectra. In our case, *GFP2* as part of bound *HAC1*^NRE^-GFP-nucleolin module and *YFP* as part of Ire1-*YFP* fluorophores serve as the donors and acceptor respectively. The appropriate strain harboring either all three plasmids i.e. *HAC1*^NRE^, GFP2-Nucleolin, and Ire1-YFP or only GFP2-Nucleolin were grown till the OD_600_ of the culture reaches 0.6 in the selective medium along with unlabelled cell. 1µg/mL of Tunicamycin was added to each culture and aliquots of cells were harvested at indicated times. Fluorescence intensities of each sample were determined by single laser GFP filter (FL1) of BD Verse Flow Cytometer. Mean fluorescence intensity is calculated by BD Accuri™ C6 software by taking 10,000 cells/sample. Average FRET efficiency (E) of the measured population according to the following equation:

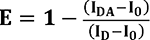

where, **I_DA_** is fluorescence intensity of donor-acceptor sample, **I_D_** is fluorescence intensity of donor only sample, and **I_0_** is fluorescence intensity of unlabeled sample.

#### Statistical Analyses

All the quantitative experiments reported in this paper (mRNA steady-state levels, decay rates, chromatin immunoprecipitation, RNA immunoprecipitation experiments, quantitative growth experiments in liquid media) were done from at least three independent sample size (biological replicates) (n=3). However, for calculating co-localization Index, the sample sizes are six (n=6). For each biological replicate, a given yeast strain was grown and treated under the same conditions independently before a given experiment was conducted. Technical replicate, in contrast, involved repetition/analyses of the same biological replicate sample for many times. A technical replicate was used to establish the variability (experimental error) involved the analysis, thus allowing one to set the confidence limits for what is significant data. All the statistical parameters such as mean, standard deviations, P-values, standard error of the mean were estimated using Graphpad Prism version 8.0.1. P-values were determined using paired Student’s two-tailed t-test.

## Supporting information

PairaS22 Bio Res Art Supporting Information

## DATA AVAILABILITY

All the raw data leading to establishing the final results presented here are available in the supplementary material as a separate excel file ‘**PairaS22 JBC Res Art Manuscript Raw Data**’

## SUPPORTING INFORMATION

This article contains supporting information that describes the list of yeast strains, plasmids, oligonucleotides, antibodies and other reagents and their sources used in this investigation. The supporting information section also contains all the raw data associated with main figures that are presented either in the form of supplementary tables and an excel file. List of references cited in the supporting information follow the list references cited in the main manuscript.

## Abbreviations

CTEXT: <u>C</u>bc1p-<u>T</u>if4631p-dependent <u>Ex</u>osome <u>T</u>argeting
UPR: <u>U</u>nfolded <u>P</u>rotein <u>R</u>esponse
BE: <u>B</u>ipartite <u>E</u>lement
ER: <u>E</u>ndoplasmic <u>R</u>eticulum,

## ACKNOWLEDGMENTS

We gratefully acknowledge Dr. Randy W. Schekman (University of California, Berkeley) for kindly sharing the *ypt1-3* mutant yeast strain and Dr. Stephen Buratowski (Harvard Medical School, Boston, MA, USA) for sharing *nrd1-1* mutant strain, and anti-Nrd1 antibody. We also acknowledge Dr. Scott Butler for providing us with the Anti-Rrp6p, Anti-Rrp4p, Anti-Cbc1p, and Anti-Tif4631 antibodies (University of Rochester, Rochester, NY, USA). We thank Dr. Satarupa Das, and the members of the Das Laboratory for critically reading this manuscript. We also thank the anonymous reviewers for their critical comments and constructive suggestions.

## AUTHOR CONTRIBUTIONS

SP and AC conducted all the experiments. SP and BD conceived and designed all the experiments, BD directed the research program, and organized and wrote the manuscript.

