## Supplementary material for "The Rab GTPase Ypt1p governs the activation of Unfolded Protein Response (UPR) in *Saccharomyces cerevisiae* by promoting the preferential nuclear degradation of pre-*HAC1* mRNA": PairaS22 Bio Res Art Supporting Information

**RESEARCH ARTICLE** **Submitted to *J. Biol. Chem.***

### The Rab GTPase Ypt1p controls the activation of Unfolded Protein Response (UPR) in *Saccharomyces cerevisiae*by facilitating the recruitment of the NNS complex, and the nuclear exosome/CTEXT components to pre-*HAC1* mRNA

Sunirmal Paira, Anish Chakraborty & Biswadip Das*

Department of Life Science and Biotechnology, Jadavpur University, Kolkata, India.

### SUPPORTING INFORMATION

**Lists of materials Included in the supporting information**

1. **Supplementary Table S1: List of yeast strains used in this study**
2. **Supplementary Table S2: List of plasmids used in this study**
3. **Supplementary Table S3: List of primers used in this study**
4. **Supplementary Table S4: List of Antibodies used in this study and their sources**
5. **Supplementary Table S5: Co-localization index of different strains (Data associated with Fig. 4B)**
6. **Supplementary Table S6: Efficiencies of FRET in WT and *ypt1-3* yeast strains (Data associated with Fig. 4C)**
7. **Supplementary Table S7: Co-localization index in WT and *ypt1-3* yeast strains (Data associated with Fig. 6C and D)**
8. **Supplementary Figure S1 with Legend**
9. **Supplementary Figure S2 with Legend**
10. **Supplementary Figure S3 with Legend**
11. **Supplementary Figure S4 with Legend**

### SUPPLEMETARY TABLES

#### Supplementary Table S1: List of yeast strains used in this study

| Strain | Genotype | Reference |
|---|---|---|
| yBD117 | *Mata leu2-3,112 ura3-52 his3-Δ200 trp1-Δ901 lys2-801* | (1) |
| yBD118 | *Mata leu2-3,112 ura3-52 his3-Δ200 trp1- Δ901 lys2-801 hac1::TRP1* | (1) |
| yBD129 | *Mata leu2-3,112 ura3-52 his3- Δ200 trp1- Δ901 lys2-801 rrp6::URA3* | (2) |
| yBD131 | *Mataleu2-3,112 ura3-52 his3- Δ200 trp1- Δ901 lys2-801 cbc1:: hisG* | (2) |
| yBD148 | *MATa leu2-3,112 ura3-52 trp1-1 his3-11,15 nrd1Δ::HIS3; lys2Δ2; ade2-1; met2Δ1, can1-100 [pRS316NRD1 (NRD1, URA3, CEN)]* | (3) |
| yBD157 | *MATα ura3Δ0 his3Δ1 leu2Δ0; met15Δ0 LYS2 nrd1Δ::KAN RRP6-TAP::HIS3 [pJC580 (NRD1-HA, LEU2, CEN)]* | (3) |
| yBD158 | *MATα ura3Δ0 his3Δ1 leu2Δ0; met15Δ0; LYS2; nrd1Δ::KAN; RRP6-TAP::HIS3 [pJC720 (nrd1-102-HA, LEU2, CEN)]* | (3) |
| yBD177 | *MATa ura3-52 leu2-3,112 trp1-1 his3-11,15 nrd1::HIS3 lys2Δ2 ade2-1 met2Δ1 can1-100[pRS 424nrd1-1(nrd1-1, TRP1, 2μ ori)]* | (3) |
| yBD276 | *Mataura3-52 leu2-3,112 his3-Δ200 trp1-Δ901 lys2-801 HIS3-GAL::protA-CBC1 (pGAL-cbc1)* | (2) |
| yBD433 | *MATa ura3-52 leu2-3,112* | (4) |
| yBD434 | *MATa ura3-52 leu2-3,112ypt1-3* | (4) |
| yBD457 | *MATa ura3-52 leu2-3,112 YPT1-TAP:: URA3* | This Study |
| yBD549 | *Mata leu2-3,112 ura3-52 his3-Δ200 trp1-Δ901 lys2-801hac1::TRP1 [pRS316 HAC1(HAC1,URA3, CEN)]* | This Study |
| yBD550 | *Mataleu2-3,112 ura3-52 his3-Δ200 trp1-Δ901 lys2-801hac1::TRP1 [pRS316 HAC1-ΔR1(HAC1- ΔR1,URA3,CEN)]* | This Study |
| yBD551 | *Mata leu2-3,112 ura3-52 his3-Δ200 trp1-Δ901 lys2-801 hac1::TRP1 [pRS316 HAC1-ΔR2(HAC1- ΔR2,URA3,CEN)]* | This Study |
| yBD552 | *Mata leu2-3,112 ura3-52 his3-Δ200 trp1-Δ901 lys2-801 hac1::TRP1 [pRS316 HAC1 ΔBE(HAC1- ΔBE,URA3,CEN)]* | This Study |
| yBD553 | *Mata leu2-3,112 ura3-52 his3-Δ200 trp1-Δ901 lys2-801hac1::TRP1 [pRS316HAC1 Δ3'UTR(HAC1- Δ3'UTR,URA3,CEN)]* | This Study |
| yBD565 | *MATa ura3-52 ypt1-3-TAP::URA3* | This Study |
| yBD572 | *MATa ura3-52 leu2-3,112 trp1-1 his3-11,15 nrd1::HIS3 lys2Δ2 ade2-1 met2Δ1 can1-100 [pRS316NRD1 (NRD1, URA3, CEN), pRS 315 YPT1-TAP(YPT1-TAP,LEU2,CEN)]* | This Study |
| yBD573 | *MATa ura3-52 leu2-3,112 trp1-1 his3-11,15 nrd1::HIS3 lys2Δ2 ade2-1 met2Δ1 can1-100, [pRS 424nrd1-1(nrd1-1, TRP1, 2μ ori), pRS 315 YPT1-TAP(YPT1-TAP,LEU2,CEN)]* | This Study |
| yBD579 | *MATa ura3-52 ypt1-3 hac1::hisG [pRS316 HAC1-ΔR1(HAC1- ΔR1,URA3,CEN)]* | This Study |
| yBD580 | *MATa ura3-52 ypt1-3 hac1::hisG [pRS316 HAC1-ΔR2(HAC1- ΔR2,URA3,CEN)]* | This Study |

| | | |
|---|---|---|
| yBD581 | *MATa ura3-52 ypt1-3 hac1::hisG [pRS316HAC1 Δ3'UTR(HAC1- Δ3'UTR,URA3,CEN)]* | This Study |
| yBD582 | *MATa ura3-52 ypt1-3 hac1::hisG [pRS316HAC1 Δ3'UTR(HAC1- ΔBE,URA3,CEN)]* | This Study |
| yBD583 | *Mata leu2-3,112 ura3-52 his3-Δ200 trp1-Δ901 lys2-801 hac1::TRP1 cbc1::hisG [pRS316 HAC1-ΔR1(HAC1- ΔR1,URA3,CEN)]* | This Study |
| yBD584 | *Mataleu2-3,112 ura3-52 his3-Δ200 trp1-Δ901 lys2-801 hac1::TRP1 cbc1::hisG [pRS316 HAC1-ΔR2(HAC1- ΔR2,URA3,CEN)]* | This Study |
| yBD585 | *Mataleu2-3,112 ura3-52 his3-Δ200 trp1-Δ901 lys2-801 hac1::TRP1 cbc1::hisG [pRS316 HAC1 ΔBE(HAC1- ΔBE,URA3,CEN)]* | This Study |
| yBD588 | *Mata leu2-3,112 ura3-52 his3-Δ200 trp1-Δ901 lys2-801 hac1::TRP1 [pRS 315 YPT1-TAP(YPT1-TAP,LEU2,CEN), pRS316 HAC1(HAC1,URA3, CEN)]* | This Study |
| yBD589 | *Mata leu2-3,112 ura3-52 his3-Δ200 trp1-Δ901 lys2-801hac1::TRP1[pRS 315 YPT1-TAP(YPT1-TAP,LEU2,CEN), pRS316 HAC1-ΔR1(HAC1- ΔR1,URA3,CEN)]* | This Study |
| yBD590 | *Mata leu2-3,112 ura3-52 his3-Δ200 trp1-Δ901 lys2-801hac1::TRP1[pRS 315 YPT1-TAP(YPT1-TAP,LEU2,CEN), pRS316 HAC1-ΔR2(HAC1- ΔR2,URA3,CEN)]* | This Study |
| yBD591 | *Mata leu2-3,112 ura3-52 his3-D 200 trp1-Δ901 lys2-801hac1::TRP1[ pRS 315 YPT1-TAP(YPT1-TAP,LEU2,CEN), pRS316 HAC1-ΔBE(HAC1- ΔBE,URA3,CEN)]* | This Study |
| yBD592 | *Mata leu2-3,112 ura3-52 his3-D 200 trp1-Δ901 lys2-801 hac1::TRP [pRS 315 YPT1-TAP(YPT1-TAP,LEU2,CEN), pRS316 HAC1- Δ3'UTR(HAC1- Δ3'UTR,URA3,CEN)]* | This Study |

**Supplementary Table S2: List of plasmids used in this study**

| Plasmid | Genotype | Reference |
|---|---|---|
| pBD42 | *CBC1* disruptor, *pUC18 cbc1::URA3* | (5) |
| pBD217 | *ND-GFP2* in low-copy *LEU2* | (6) |
| pBD218 | *Ire1-RFP in high-copy LEU2 vector* | (6) |
| pBD219 | *HAC1*$^{NRE}$*in pRS 316 URA3 vector* | (6) |
| pBD93 | *HAC1 disrupter; pRS 316 hac1::URA* | This study |
| pBD95 | *HAC1* inserted in the *Bam*HI site of *pRS 316* | This study |
| pBD220 | *HAC1*-ΔBE *pRS 316 CEN* | (6) |
| pBD333 | 154 bp of 3′-UTR of *HAC1* was deleted from *pRS 316HAC1* by reverse PCR using primer pair OBD 842 & OBD843 | This study |
| pBD334 | 142 bp of 3′-UTR of *HAC1*was deleted from *pRS 316HAC1* by reverse PCR using primer pair OBD 844 & OBD845. | This study |
| pBD335 | 379 bp of *HAC1* 3′UTR (1-379) was deleted from *pRS 316HAC1* by reverse PCR using primer pair OBD 842 & OBD 845. | This study |
| pBD342 | Ypt1-TAP tagged gene cloned in Sma1 site of *pRS 315 LEU2 CEN* | This study |

**Supplementary Table S3: List of primers used in this study**

| Primer | Oligonucleotide Sequence | Gene to be amplified |
|---|---|---|
| OBD321<br>OBD322 | Forward 5′-GCGGGAAACAGTCTACCCTTT-3′<br>Reverse 5′-CACTGTAGTTTCCTGGTCATCGTAA-3′ | Pre-*HAC1* mRNA (intron specific primer) |
| OBD325<br>OBD326 | Forward 5′-ATTGACACTGCTCGTTTGTGTGT-3′<br>Reverse 5′-CGAGACGACAATGGGATTGA-3′ | *IRE1* mRNA |
| OBD323<br>OBD324 | Forward 5′-TTTCCTGAACAAATAGAGCCATTCT-3′<br>Reverse 5′-TGCGCTTCGGACAGTACAAG-3′ | *HAC1* Intron-Exon 2 |
| OBD522<br>OBD523 | Forward 5′-ACATCGTACCGAGTGATGAACG-3′<br>Reverse 5′-GGCGGTTGTTGTCGTAGGTG-3′ | *HAC1* Promoter |
| OBD537<br>OBD539 | Forward 5′-CGCAATCGAACTTGGCTATCCCTACC 3′<br>Reverse 5′-CCCACCAACAGCGATAATAACGAG 3′ | Pre- and Mature *HAC1* mRNA |
| OBD184<br>OBD185 | Forward 5′-AAAGGGTGGCCCACATAAGG- 3′<br>Reverse5′-CTTCAACCCACCAAAGGCCA-3′ | *CYC1* |
| OBD272<br>OBD273 | Reverse 5′-GTGTGGATTTGATGGTATGTGTGA-3′<br>Forward 5′-GATATGCAGGGTCGATAACTGAAA-3′ | *LYS2* |
| OBD276<br>OBD277 | Forward 5′-AGCGTCCGTAATGTCCAATTCT-3′<br>Reverse 5′-CCCGCACCATAAGCTATGTGA-3′ | *NCW2* |
| OBD607<br><br><br>OBD608 | Forward5′-<br>GACAAAGGGAACGTCAACCTGAAGGGACAGAGTTTAACCAACACCGGTGG<br>GGGCTGCTGTTCCATGGAAAAGAGAAG-3′<br>Reverse 5′-<br>GTCTTATTTACTTATTTAGTTATTATATTATATGGGTCTGCAAGGTAGAGGC<br>GCGCTTGTTACGACTCACTATAGGG-3′ | Tap tagging of *YPT1* gene |
| OBD842<br>OBD843 | HAC1-3′ΔR1 Rev-5′-TTATACCCTCTTGCGATTGTCT-3'<br>HAC1-3′ΔR1 For-5′-CTTCAACCGAAGAAGAAGAGG-3' | Construction of *HAC1*-ΔR1 |

| | | |
|---|---|---|
| OBD844 | HAC1-ΔBE REv-5′-GCCAAAAAAGGGTCTTGTTC-3′ | Construction of *HAC1*-ΔBE |
| OBD845 | HAC1- ΔBE Rev-5′TGTTTGTGTCATATCTATCTGTC-3′ | |
| OBD846 | HAC1-Strepto 3-Δ1 F-5′-TAATACGACTCACTATAAGACAATCGCAAGAGGGTATAA-3′ | Streptotag affinity purification |
| OBD851 | HAC1-Strepto 3-Δ1 R-5′-GGATCCGACCGTGGTGCCCTTGCGGGCAGAAGTCCAAATGCGATCCCCTCTTCTTCTTCGGTTGAAG-3′ | |

**Supplementary Table S4: List of Antibodies used in this study and their sources**

| Sl. No. | Name of the Primary Antibody | Source | Primary Antibody Dilution | Secondary Antibody | Secondary Antibody Dilution |
|---|---|---|---|---|---|
| 1 | Anti-TAP | Commercially Procured, Thermo Scientific | 1:1000 | Anti- Rabbit | 1:3000 |
| 2 | Anti-Rrp6 | Dr. Scott Butler, University of Rochester | 1:1000 | Anti- Rabbit | 1:3000 |
| 3 | Anti-Cbc1 | Dr. Scott Butler, University of Rochester | 1:1000 | Anti- Rabbit | 1:3000 |
| 4 | Anti-Rrp4 | Dr. Scott Butler, University of Rochester | 1:1500 | Anti- Rabbit | 1:3000 |
| 5 | Anti-Tif4631 | Dr. Julie Baron-Benhamou, Rockefeller University, New York | 1:2000 | Anti- Rabbit | 1:3000 |
| 6 | Anti-Nrd1 | Dr. David Brow, University of Wisconsin Madison | 1:1000 | Anti- Rabbit | 1:3000 |
| 7 | Anti-Nab3 | Dr. Maurice S. Swanson, University of Florida | 1:1000 | Anti-Mouse | 1:3000 |

**Supplementary Table S5: Co-localization index of different strains (Data associated with Fig. 4B)**

| Sample | Co-localizationIndex | |
|---|---|---|
| | WT | *ypt1-3* |
| 1 | 0.2314 | 0.7891 |
| 2 | 0.2765 | 0.6792 |
| 3 | 0.3532 | 0.7623 |
| 4 | 0.4213 | 0.4543 |
| 5 | 0.2341 | 0.7832 |
| 6 | 0.2156 | 0.4698 |
| 7 | 0.2451 | 0.5981 |
| 8 | 0.2561 | 0.6923 |
| **Mean** | **0.2791** | **0.6535** |

**Supplementary Table S6: Efficiencies of FRET in WT and *ypt1-3* yeast strains (Data associated with Fig. 4C)**

| FRET efficiencies in WT cells | | | | |
|---|---|---|---|---|
| **Donor-Acceptor ($HAC1^{NRE}$-GFP2+Ire1p-RFP) $I_{DA}$** | **Only Donor($HAC1^{NRE}$-GFP2) $I_D$** | **Unlabeled sample $I_0$** | **FRET efficiency, $E = 1 - \frac{(I_{DA}-I_0)}{(I_D-I_0)}$** | **Mean FRET WT** |
| 571 | 742 | 524 | 0.7251 | |
| 517 | 793 | 410 | 0.7206 | **0.6575** |
| 621 | 738 | 516 | 0.5270 | |

| FRET efficiencies in *ypt1-3* cells | | | | |
|---|---|---|---|---|
| **Donor-Acceptor ($HAC1^{NRE}$-GFP2+Ire1p-RFP) $I_{DA}$** | **Only Donor($HAC1^{NRE}$-GFP2) $I_D$** | **Unlabeled sample $I_0$** | **FRET efficiency, $E = 1 - \frac{(I_{DA}-I_0)}{(I_D-I_0)}$** | **Mean FRET *ypt1-3*** |
| 658 | 991 | 682 | 1.0776 | |
| 662 | 1194 | 671 | 1.0172 | **1.0848** |
| 621 | 897 | 659 | 1.1596 | |

**Supplementary Table S7: Co-localization index in WT and *ypt1-3* yeast strains (Data associated with Fig. 6C and D)**

| Co-localization Index of WT Ypt1p and Nuclear signals | | |
|---|---|---|
| **Ypt1p-TAP** | **Stressed** | **Unstressed** |
| 1 | 0.185 | 0.4778 |
| 2 | 0.2996 | 0.6048 |
| 3 | 0.2817 | 0.427 |
| 4 | 0.3096 | 0.4391 |
| 5 | 0.3564 | 0.3718 |
| 6 | 0.2165 | 0.5956 |
| **Mean** | **0.2748** | **0 .4860** |

| Co-localization Index of mutant Ypt1-3p and Nuclear signals | | |
|---|---|---|
| **Ypt1-3p-TAP** | **Stressed** | **Unstressed** |
| 1 | 0.4056 | 0.4232 |
| 2 | 0.3714 | 0.3849 |
| 3 | 0.2985 | 0.3567 |
| 4 | 0.4231 | 0.3876 |
| 5 | 0.3678 | 0.3276 |
| 6 | 0.5346 | 0.5005 |
| **Mean** | **0.4002** | **0.3968** |

### SUPPLEMETARY FIGUERES WITH LEGENDS

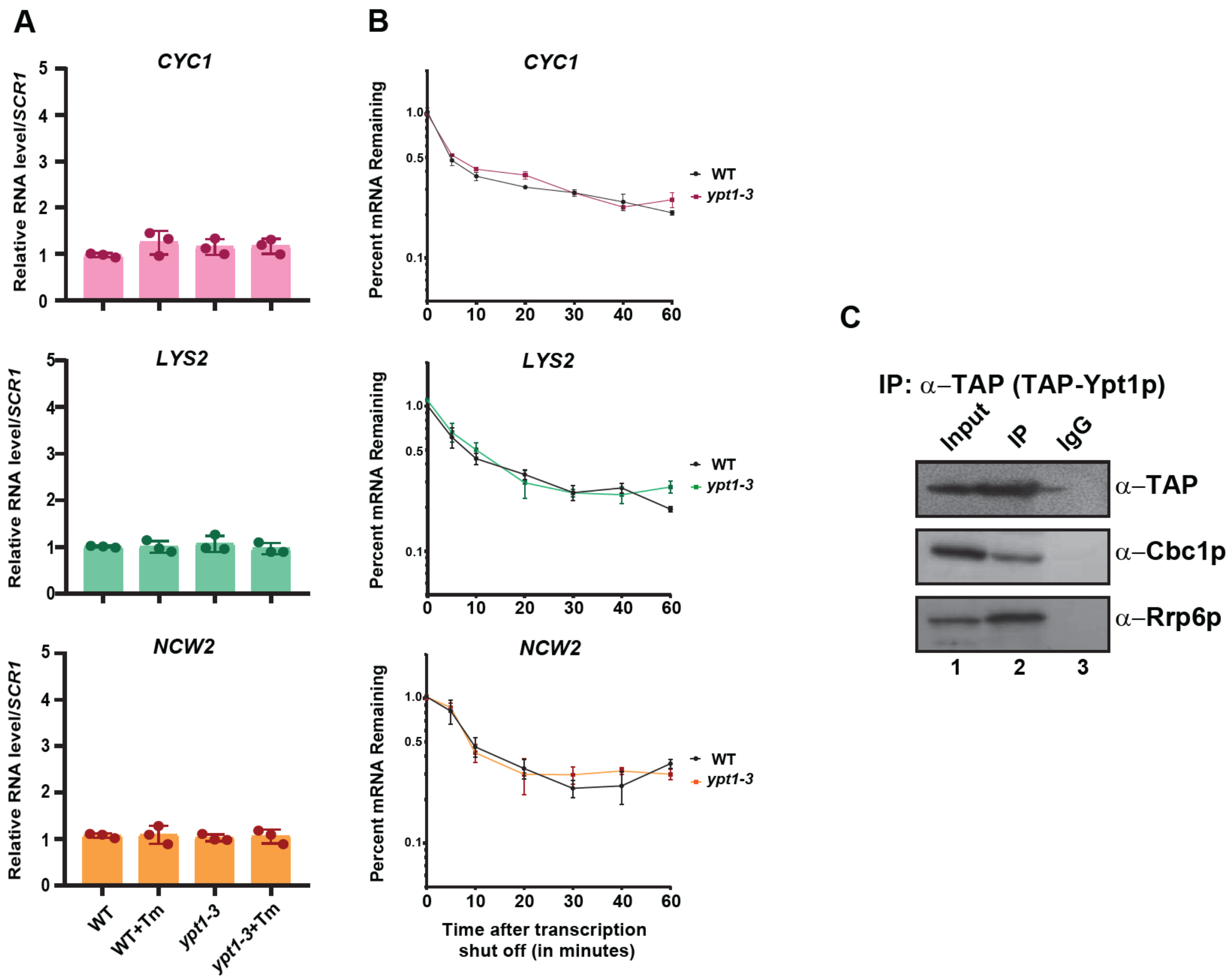


**Figure S1:** **A.** Histogram depicting the steady state levels of *CYC1*, *LYS2,* and *NCW2*mRNA levels in WT (yBD-433), *ypt1-3* (yBD-434) yeast strains in the absence and presence of tunicamycin as determined by qPCR analysis. Transcript copy numbers/2ng cDNA of each strain was normalized to *SCR1*RNA levels in respective strains and are presented as means ± SE (n=3 biological replicates for each strain). Normalized value of individual mRNA from WTstrain was set to one. The statistical significance of difference as reflected in the ranges of p-values estimated from Student's two-tailed t-tests for a given pair of test strains for each message are presented with following symbols, ∗<0.05, ∗∗<0.005, and ∗∗∗<0.001, NS, not significant. **B.**Decay rates of *CYC1*, *LYS2* and *NCW2* mRNA in WT and *ypt1-3* strains. Decay rates were determined from three independent experiments (biological replicates) by qRT-PCR analysis (using primer sets corresponding to *HAC1* intronic sequence) and the intronic signals were normalized to *SCR1* RNA and normalized signals (mean values ± SD) were presented as the fraction of remaining RNA (with respect to normalized signals at 0 min) vs. time of incubation in the presence of 1, 10-phenanthroline. **C.** Ypt1p strongly interacts with the CTEXT component Cbc1p and exosome component Rrp6p in the absence of ER stress. Lane1 detection of Ypt1p, Cbc1p and Rrp6p, using their respective antibodies from their input samples. Lane 2 detection of Ypt1p, Cbc1p and Rrp6p from their immunoprecipitate samples and Lane 3 Mock IP using IgG

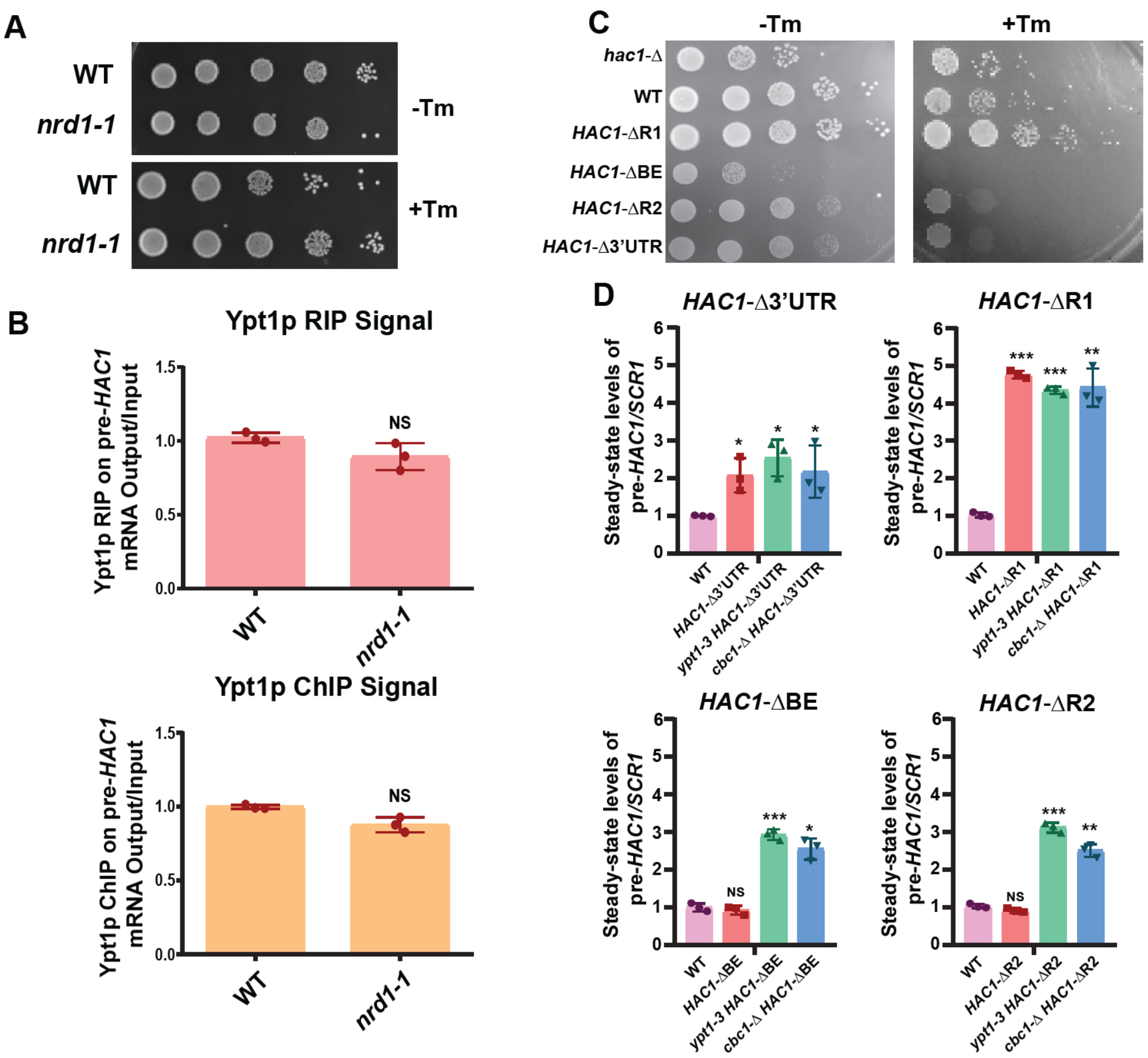


**Figure S2 A.** Relative growth of WT and mutant *nrd1-1* yeast strains in YPD solid growth media in absence (-Tm) and in presence of 1µg/ml tunicamycin (+Tm). Equal no. of cells of these yeast strains grown in YPD liquid medium in absence of tunicamycin were spotted with ten-fold dilutions on medium with or without tunicamycin and grown at 30°C for 72 hours before photographed. **B-C.** Relative binding affinity (**B**) and co-transcriptional recruitment (**C**) of Ypt1p on to precursor *HAC1* message in WT and *nrd1-1* strain. Extracts were prepared from WT and *nrd1-1*

yeast cells expressing Ypt1p-TAP following RNA-protein or chromatin-protein crosslinking as described before and was subjected to RIP or Chromatin-IP. Quantification of the bound pre-HAC1 RNA was then estimated. Normalized value of individual pre-*HAC1* RNA recovered from the RIP/ChIP sample from WT strain was set to one. The statistical significance of difference as reflected in the ranges of p-values estimated from Student's two-tailed t-tests for a given pair of test strains for each message are presented with following symbols, ∗<0.05, ∗∗<0.005, and ∗∗∗<0.001, NS, not significant. D**.** Relative growth of *HAC1*-FL, *HAC1*-Δ3'UTR, *HAC1*-ΔR1, *HAC1*-ΔBE and *HAC1*-ΔR2 strains in YPD solid growth media in absence (-Tm) and in presence of 1μg/ml tunicamycin (+Tm). Equal no. of cells of these yeast strains grown in YPD liquid medium in absence of tunicamycin were spotted with ten-fold dilution on medium with or without tunicamycin and grown at 30°C for 72 hours and photographed. **E.** The relative steady state levels of full length *HAC1* mRNA and its various 3′-UTR deleted versions in WT, *ypt1-3*, *cbc1*-Δ strains using qRT-PCR analysis with a primer pair specific to *HAC1*intron. Transcript copy numbers/2 ng cDNA of each strain was normalized to *SCR1*RNA levels in respective strains and are presented as means ± SE (n=3 for each strain). Normalized value of individual mRNA from normal samples was set to one. The statistical significance of difference as reflected in the ranges of p-values estimated from Student's two-tailed t-tests for a given pair of test strains for each message are presented with following symbols, ∗<0.05, ∗∗<0.005 and ∗∗∗<0.001, NS, not significant.

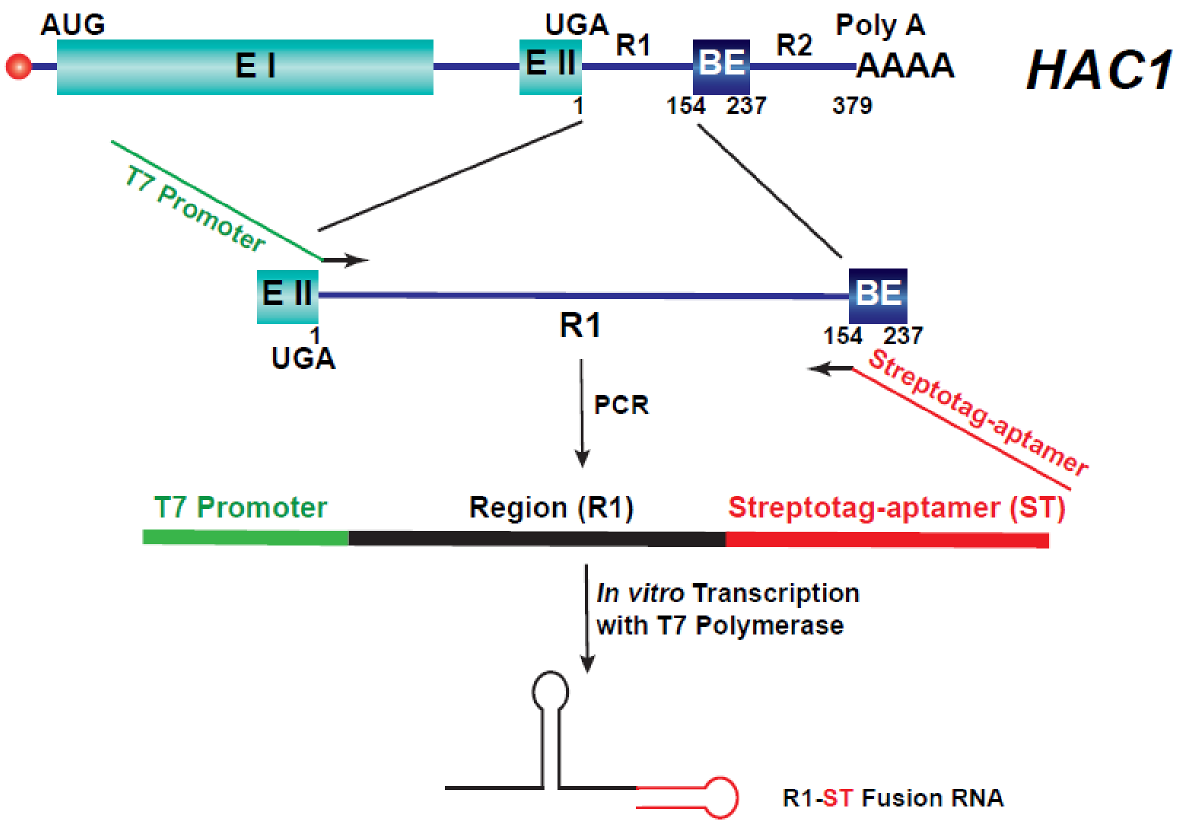


**Figure S3: Strategy to prepare HAC1-R1-Streptotag (R1-ST) aptamer fusion RNA**. Cloned version of *HAC1* gene was used as a template for a PCR reaction using a set of overhang primer sets. The forward primer harbors a sequence corresponding to functional T7 promoter (indicated by green line) followed by a sequence complementary to the 5′-end of the region 1 (R1) of the *HAC1* 3′-UTR (indicated by a black line with right arrowhead). The reverse primer harbors a sequence corresponding to Streptotag aptamer (indicated by red line) followed by a complementary sequence corresponding to the 3′-end of the region 1 (R1) of the *HAC1* 3′-UTR (indicated by a black line with right arrowhead). The product that yielded from such reaction contains sequences corresponding to T7 promoter (Green), *HAC1* 3′-UTR region 1 (R1) (black), and Streptotag aptamer (red) in tandem. This linear DNA was used in *in vitro* transcription reaction using T7 RNA polymerase to produce copious amount of R1-ST fusion RNA that was bound to the dihydrostreptomycin coupled to epoxy-activated sephadex-6B.

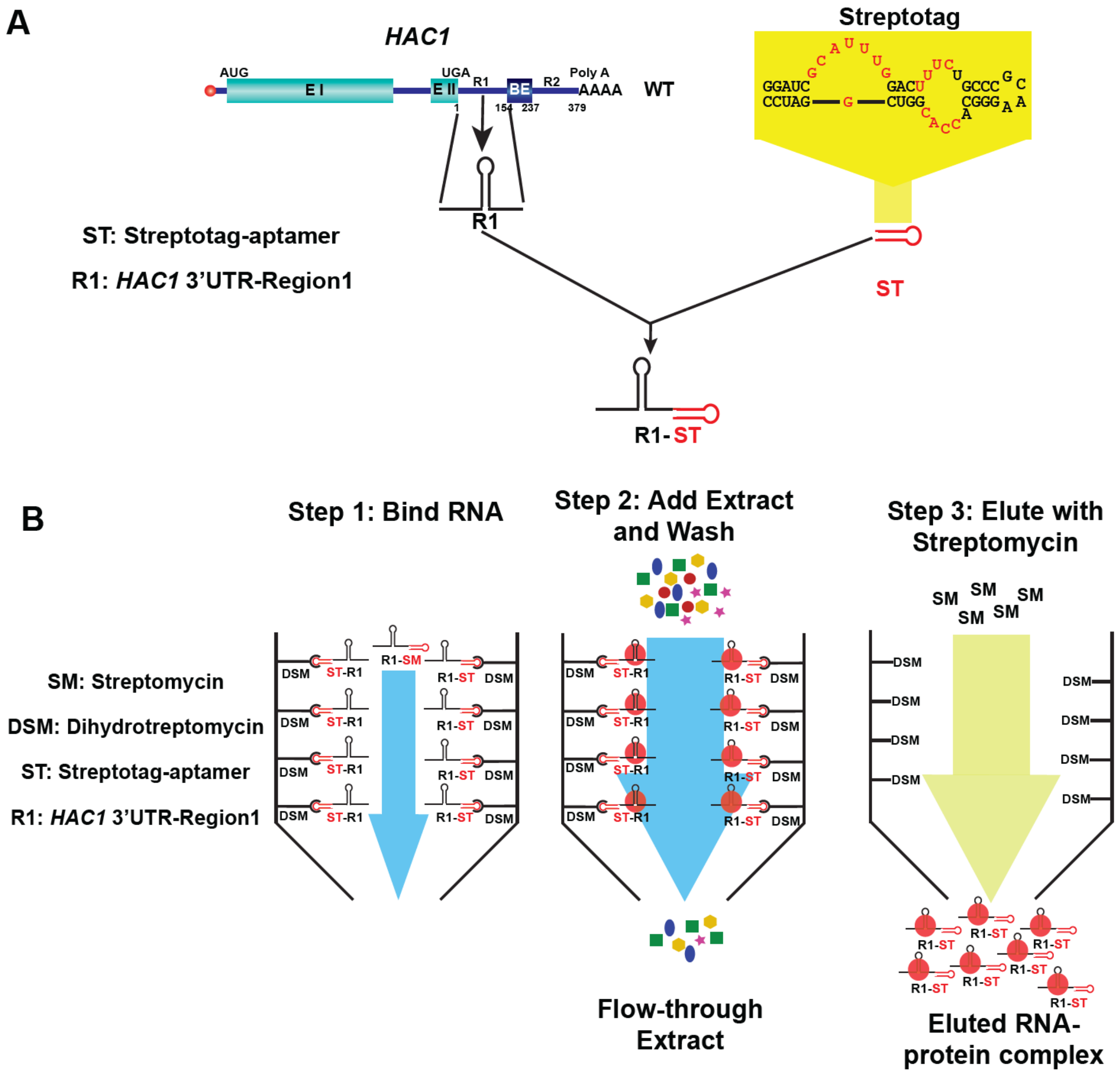


**Figure S4:** A. Schematic diagram showing the structure of the *HAC1* R1-ST fusion RNA. B. Cartoon representing the rationale of the Streptotag affinity purification procedure depicting three major sequential steps involved. The bait R1-ST fusion RNA is first bound to dihydrostreptomycin (DSM) coupled to epoxy-activated sephadex-6B followed by the addition of the cellular or nuclear extracts from appropriate yeast strain that allows binding and immobilization of the prey protein(s). After repeated washing to get rid of the non-specific proteins, the bound RNA protein complex is eluted with streptomycin (SM), which has a higher affinity to Streptotag aptamer than dihydrostreptomycin (DSM).
